# Proprotein convertase subtilisin kexin type 9 (PCSK9) inhibition attenuates abdominal aortic aneurysm formation via enhanced macrophage-dependent efferocytosis

**DOI:** 10.64898/2026.06.22.733861

**Authors:** Michael J. Fassler, Aravinthan Adithan, Jeff Arni C. Valisno, Jonathan R. Krebs, Chelsea Viscardi, Griffin Stinson, Gwendolyn S. Gillies, Walker R. Ueland, Dan Neal, Gang Su, Shiven Sharma, Pankaj K. Singh, Ramon Sun, Matthew Gentry, Ashish K. Sharma, Gilbert R. Upchurch

**Author notes:** **Address for Correspondence:** Ashish K. Sharma, MBBS, PhD Gilbert R. Upchurch Jr., MD, DHA Department of Surgery. University of Florida PO Box 100128, Gainesville, FL 32610.

## Abstract

Abdominal aortic aneurysms (AAAs) occur predominantly in the elderly population and currently there is no effective pharmacological therapy for mitigating AAA growth and preventing impending rupture. Proprotein subtilisin kexin type 9 (PCSK9) gene has been identified as a specific risk-locus for AAA development. However, the mechanistic and clinical role of PCSK9-mediated signaling in AAAs has not been delineated. We demonstrate that treatment with PCSK9 inhibitors, such as Evolocumab, mitigates vascular inflammation and remodeling, resulting in attenuated aneurysm growth in clinical datasets as well as experimental models of AAA and aortic rupture. Mechanistically, Evolocumab immunomodulates macrophage reprogramming to enhance clearance of apoptotic smooth muscle cells via MerTK-dependent efferocytosis that ameliorates aortic inflammation and vascular remodeling. Furthermore, Evolocumab increases the expression of oxidized phosphatidylserine species and decreases expression of lysophospholipids, succinate, and glycolytic intermediates within the aortic wall compared to untreated controls, further enhancing the pro-resolving functions of macrophages. Collectively, our data demonstrates the ability of PCSK9 inhibition to regulate macrophage-specific efferocytosis that limits AAA progression and prevents aortic rupture.

## Introduction

Abdominal aortic aneurysm (AAA) is a localized dilation of the abdominal aorta, defined as a diameter exceeding 3 cm or 50% larger than a normal adjacent aortic segment.^1,2^ Established risk factors for AAA development include elderly age, male sex, and a history of smoking.^3^ Importantly, there are no pharmacologic modalities to mitigate the growth of aortic aneurysms, which can lead to sudden rupture, carrying an overall mortality of 80-90%.^4–6^ Contemporary guideline management of AAA focuses on radiographic screening of at-risk populations, with guided imaging surveillance and invasive aortic reconstructions once a threshold of rupture risk exceeds the risk of repair.^7^ The most commonly utilized predictive factor for risk of and progression to rupture is the maximum diameter of the AAA.^7,8^

AAA pathogenesis is defined by degenerative changes to the aortic wall, characterized by smooth muscle cell (SMC) apoptosis, extracellular matrix (ECM) degradation, and infiltration of inflammatory cells, such as macrophages and neutrophils.^9–11^ ECM breakdown products, accumulating cell death debris, and pro-inflammatory cytokines generate a milieu with chemokines and chemoattractants, which further recruit circulating leukocytes and macrophages into the aortic wall.^11,12^ Structural degradation is augmented by matrix metalloproteinase (MMP) release from both SMCs and infiltrating macrophages within the aortic wall.^11,13,14^ Recent studies have implicated that oxidized lipids are important mediators of the underlying chronic inflammatory processes of AAA formation, which can contribute to vascular cell death pathways such as apoptosis and ferroptosis.^15,16^ Clinical studies have indicated a negative correlation between lipid lowering agents, such as statins, and slower AAA growth rates and decreased hazard of undergoing repair, rupture, and death.^17–19^ However, the clinical trials involving statin therapy and long term AAA growth rates have been largely inconclusive.^20,21^

Previous genome-wide association meta-analyses have identified AAA risk loci involved with lipid metabolism, inflammation, and vascular and extracellular matrix remodeling.^22–24^ Recently, a pivotal association between elevated systemic levels of proprotein convertase subtilisin kexin type 9 (PCSK9) and increased risk of AAA development has been documented.^23^ PCSK9 is a zymogen that functions both intracellularly and extracellularly to prevent recycling of LDL receptor (LDLR), and in turn attenuates endocytosis and clearance of circulating lipoprotein complexes.^25–28^ When intracellular cholesterol levels are low, upregulation of sterol regulatory element-binding protein-2 (SREBP-2) induces transcription of pro-PCSK9.^25,27,28^ Pro-PCSK9 is then cleaved and activated, functioning intracellularly to bind LDLR shuttling it toward subsequent lysosomal degradation.^25,28^ Conversely, extracellular PCSK9 directs LDLR degradation through either directly binding and facilitating endocytosis of the receptor, or through complexing with circulating LDL or lipoprotein, which are then bound and endocytosed in an LDLR-mediated fashion with ensuing targeted degradation of the receptor.^25,28^ These findings have been augmented by the growing utilization of injectable monoclonal antibody therapies (Evolocumab and Alirocumab) directed toward PCSK9 inhibition, which have been shown to significantly lower LDL cholesterol levels and associated cardiovascular event risk.^29^ However, a comprehensive analysis of PCSK9-mediated cell-specific signaling in chronic vascular pathologies involving aortic inflammation and remodeling, such as AAA formation and aortic rupture, remains to be delineated.

In this study we identified that PCSK9 is a critical driver of macrophage reprogramming to stimulate aortic inflammation and SMC remodeling in a lipid-independent manner. Using preclinical and experimental models, we demonstrate that recombinant PCSK9 treatment exacerbates aortic inflammation, growth, and vascular remodeling during AAA formation. Conversely, administration of clinical grade PCSK9 inhibitors, such as Evolocumab and Alirocumab, attenuates aortic growth via reprogrammed macrophage phenotype and enhanced efferocytosis of apoptotic SMCs, resulting in restored vascular homeostasis and prevention of AAA formation and aortic rupture.

## Results

### Pharmacologic inhibition of PCSK9 attenuates aortic growth and confers survival advantage in human AAA patients

Comparative analyses were performed in patients with a radiographically confirmed diagnosis of AAA, and ≥2 computed tomography scans to compare AAA diameter growth (**Fig. 1A)**. Patients were stratified based on treatment with therapeutic modalities, i.e., no therapy (n=101), PCSK9 inhibitors (Evolocumab or Alirocumab, n=15), and statins (n=50). Within the statin cohort, 62% (n=31) of patients were on high-intensity statin therapy based on ACC/AHA dyslipidemia guidelines^30^, while the remaining 36% (n=18) and 2% (n=1) were on moderate-intensity and low-intensity statin therapy, respectively. Significant attenuation in aortic growth was observed in patients treated with PCSK9 inhibitors compared to no medication and statin-treated groups (33% vs. 73% and 98%, respectively; p<0.0001), respectively. Moreover, AAA stabilization (<2 mm increase or decrease in maximum diameter) was observed to be the highest in the PCSK9-treated group compared to no treatment and statin-treated groups (53% vs. 27% or 2%, respectively; p<0.0001). Importantly, PCSK9 inhibitor-treated patients experienced AAA regression (decrease in maximum diameter of >2 mm) during the study interval (13% vs. 0% each in no therapy and statin-treated groups; p=0.008). However, overall rates of mortality were decreased but not significantly changed in the PCSK9-treated group (13%) compared to no therapy (39%) or statin therapy (46%)-treated groups (p=0.08). Raw comparisons of rates of maximal diameter change between treatment groups were also performed that demonstrated that patients on PCSK9 inhibitors experienced significantly slower rates of AAA growth (−0.4±0.2 mm/month vs. no therapy: 0.2±0.007 mm/month; p<0.001 and statin-treated-groups: 0.1±0.006 mm/month; p<0.0001; **Fig. 1B**).

**Fig. 1.**
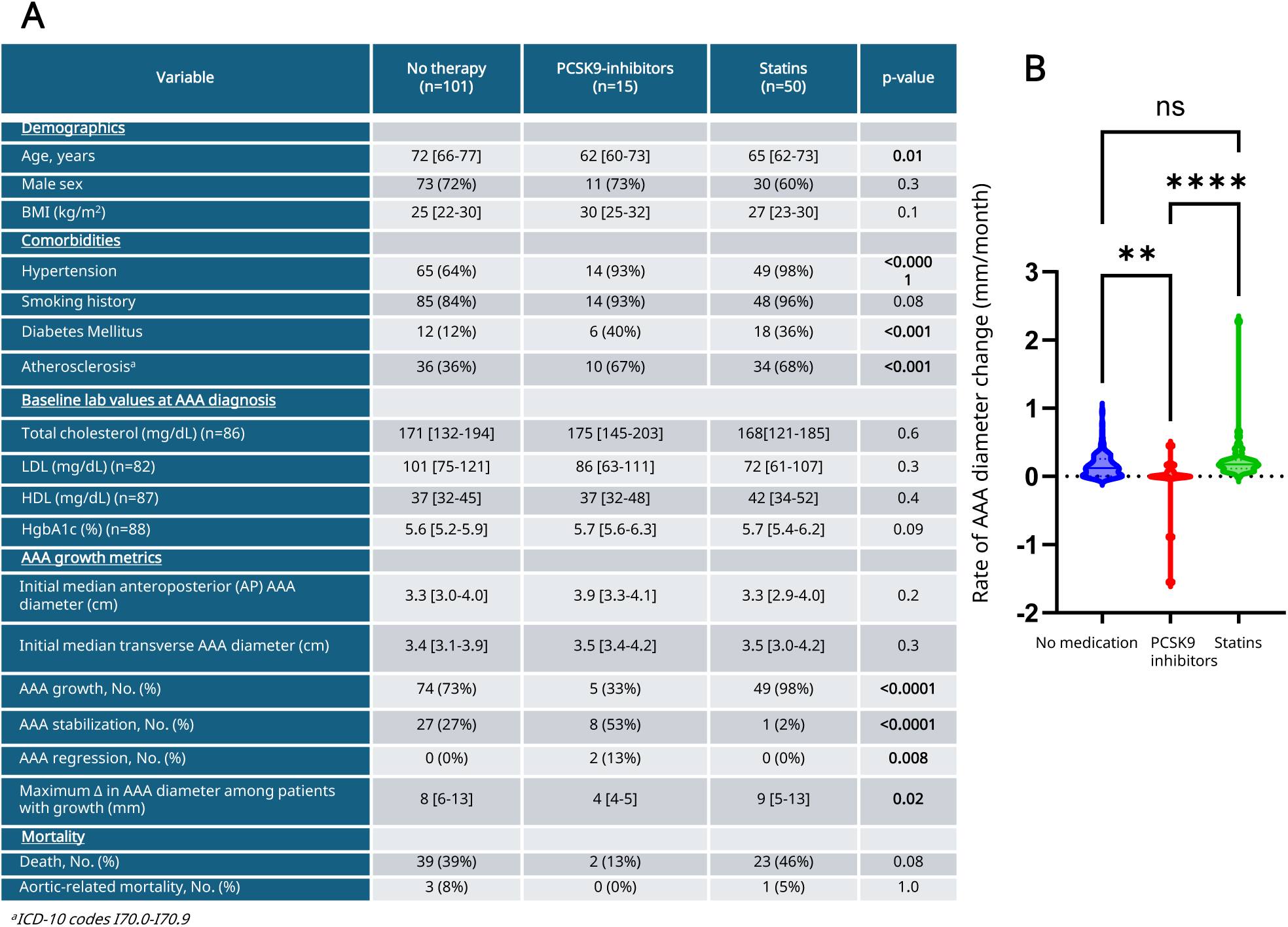
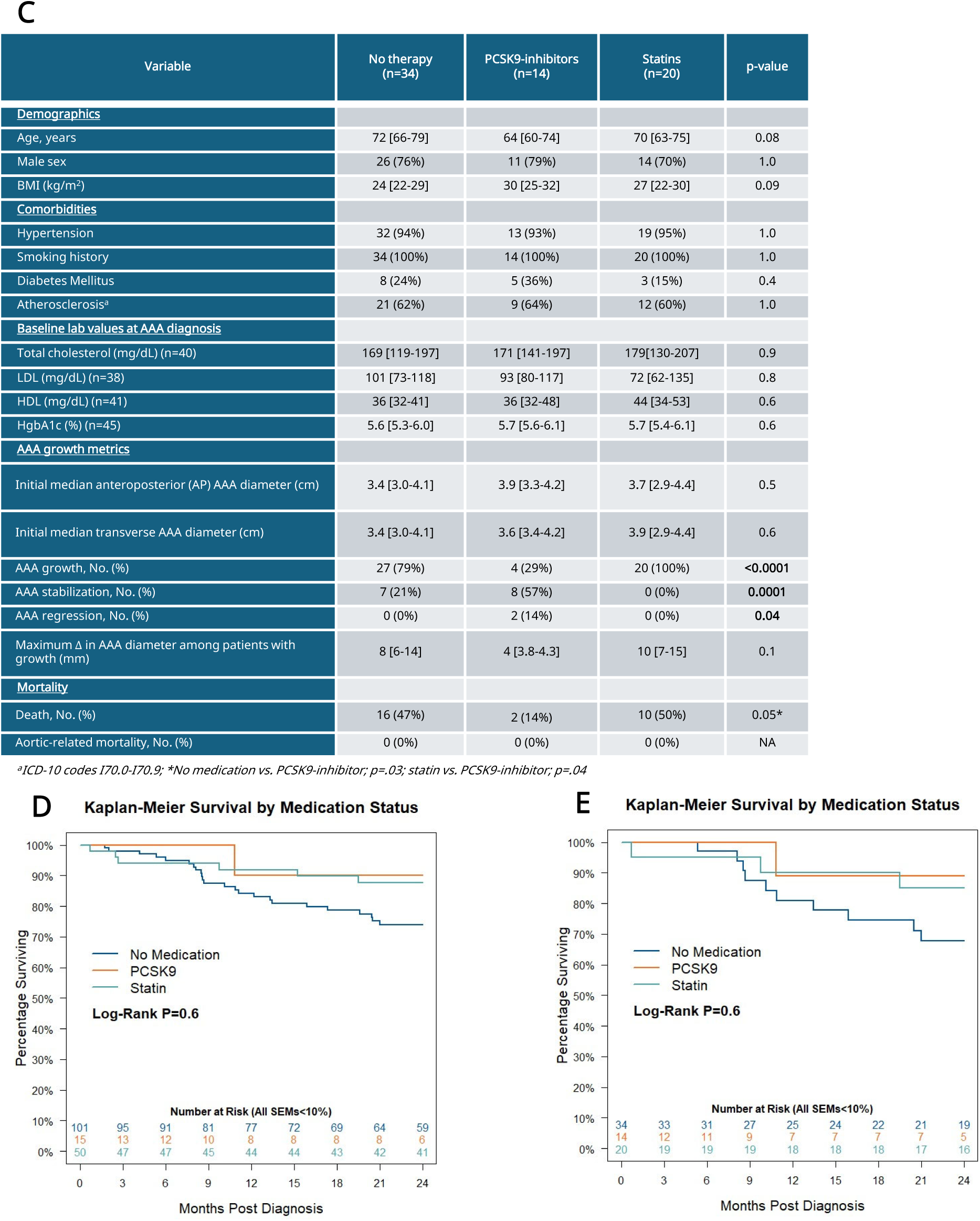
PCKS9 inhibition attenuates AAA growth and confers survival advantage in humans. **a**, Raw comparisons of clinical datasets in AAA patients stratified based on lipid therapy reveal significant differences in rates of aneurysm growth, stabilization, and regression. **b**, Raw comparisons in rates of maximum AAA diameter change (mm/month) revealed significantly decreased rates of aortic aneurysm growth with PCSK9 inhibitor therapy. **p<0.001,****p<0.0001, ns=not significant. **c**, Nearest-neighbor propensity-matched cohort comparisons (n=68) revealed significant differences in rates of AAA growth, stabilization, and regression. Additionally, pair-wise comparisons revealed significantly improved survival with PCSK9 inhibitor therapy. **d-e**, Kaplan-Meier longitudinal survival estimates between raw and propensity matched cohorts, respectively.

To correct for differences in comorbidity burden between cohorts, nearest-neighbor propensity score matching with hypertension, diabetes, smoking status, and atherosclerosis (I70.0-I70.9) as covariates was used to compare patients receiving statin or PCSK9 inhibitor therapy. Matching was performed at a ratio of 2:1, resulting in a preliminary cohort of n=27 statin patients and n=14 PCSK9 inhibitor patients. This cohort was then matched at a ratio of 1:1 on the same variables to patients receiving neither medication (n=34), patients on statin therapy (n=20), patients on PCSK9 inhibitor therapy (n=14) (**Fig. 1C**).Importantly, a, significant mitigation in the frequency of AAA growth was identified in the PCSK9 inhibitor group (29% vs. 79%; no medication vs. statin therapy; p<0.0001). This was accompanied with enhanced stabilization (57% PCSK9 inhibitor vs. 21% no medication vs. 0% statin therapy; p=0.0001), and regression (14% PCSK9 inhibitor vs. 0% no medication vs. 0% statin therapy; p=0.04). Pairwise comparisons of propensity matched cohorts revealed significant mortality benefit with PCSK9 inhibitor therapy compared to no medication (14% vs. 47%; p<0.03) and statin therapy (14% vs. 50%; p<0.05)-treated groups, respectively. Longitudinal Kaplan-Meier survival estimates were also performed for both raw cohort comparisons (**Fig. 1D**) and matched comparisons (**Fig. 1E**), demonstrating increased, but not statistically significant, survival rate at 2-years in patients treated with a PCSK9 inhibitor (raw cohort comparison: 90±9.5% PCSK9 inhibitor vs. 88±4.7% statin therapy and 74±4.6% no medication; log-rank p=0.6 and matched cohort comparison: 90±10.0% PCSK9 inhibitor vs. 85±4.7% statin therapy vs. 68±4.6% no medication; log-rank p=0.6).

### PCSK9-related genes are dysregulated in macrophages of human AAAs

To further investigate the cell specific mechanisms through which PCSK9 drives AAA formation and growth in humans, we analyzed single cell RNA sequencing data from aortic tissue from human patients with AAA and non-aneurysm control patients from two previously reported sequencing datasets (GSE226492, GSE15283).^31,32^ Differentially expressed genes in the macrophage cluster were identified and compared between patients with AAA (n=7) to non-aneurysm controls (n=5) (**Fig. 2A-B**). A total of 6277 genes were identified as variable features within the macrophage cluster and tested for differential expression (**Fig. 2C** and **Supplementary Table 1**). Of the 6277 genes, 1101 were upregulated and 929 were downregulated. 23/1101 of the upregulated and 26/929 of the downregulated genes were PCSK9-related genes (PRGs). Interestingly, literature on these 49 PRGs (**Fig. 2D-E**) highlight their role in regulating inflammation, metabolism, and phagocytosis in macrophages (STAT3, MyD88, JMJD1C, LIPIN1, SOCS3, OLR1, NLRP3, LDLR, LRP1, ANXA2, ABCA1) were dysregulated in AAAs and associated with other PCSK9 mediated disease processes.^31,33–42^ These results underscore the importance of the interaction between PCSK9 mediated signaling and macrophage phenotype during AAA pathogenesis and supported further investigation using preclinical models of AAA and aortic rupture.

**Fig. 2.**
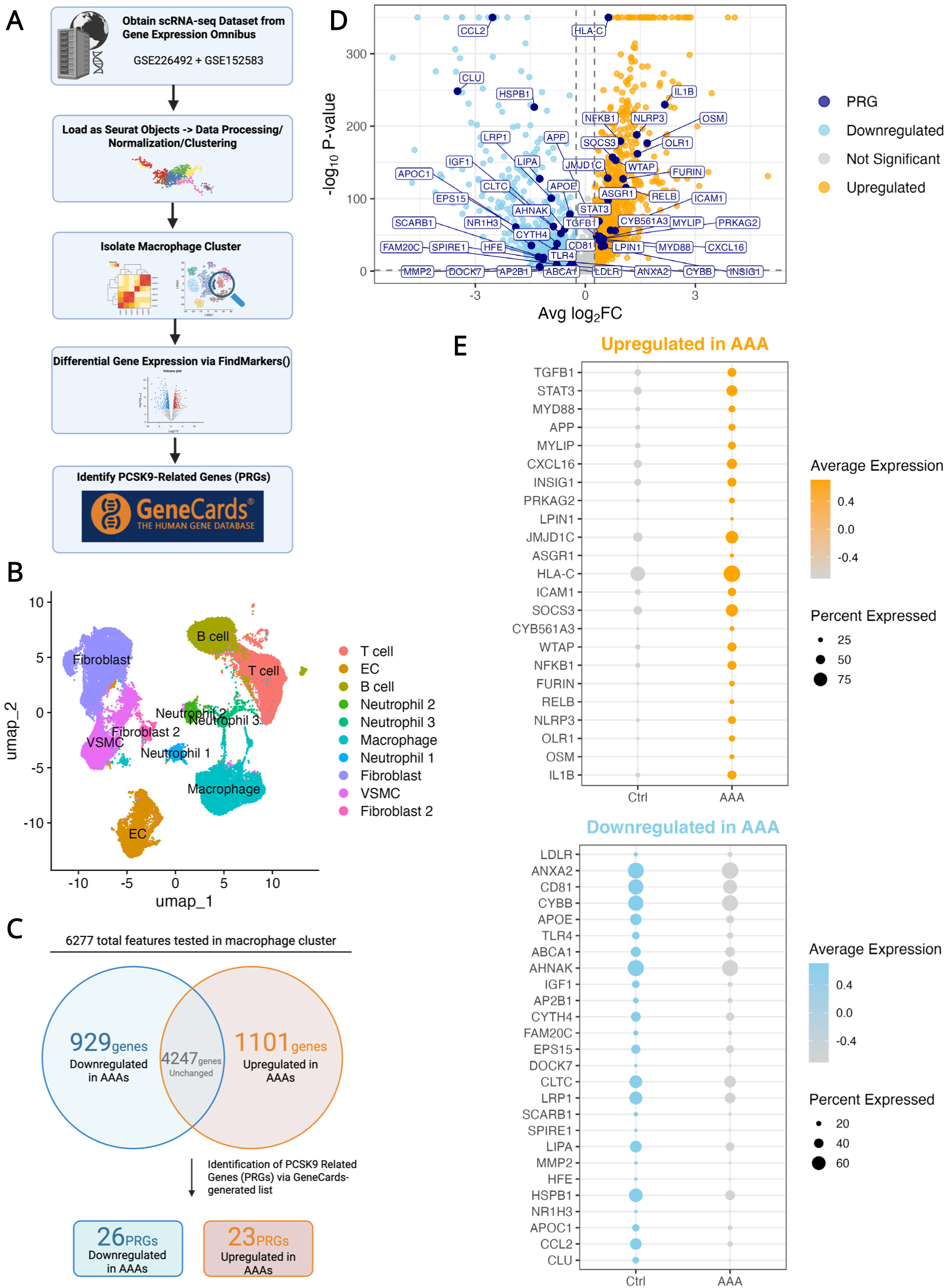
PCSK9 related genes are dysregulated in macrophages of AAA. **a**, Bioinformatic workflow to re-analyze publicly available scRNA-sequencing datasets (GSE226492, GSE152583) from Gene Expression Omnibus. Data was analyzed via Seurat and macrophage cluster was isolated. Differential expression analysis comparing macrophages from human AAAs (n=7) vs. control aortas (n=5) was performed via FindMarkers calculated using Wilcoxon testing. Genes with adjusted p-value <0.05 and |fold-change| >0.25 were considered significant differentially-expressed genes (DEGs). A list of PCSK9-related genes (PRGs) was generated using GeneCards and used to identify differentially expressed PRGs (DE PRGs). **b**, Uniform manifold projection (UMAP) plot of the annotated clusters from human AAAs and control. The macrophage cluster is highlighted and subset for further analyses. **c**, Venn diagram displaying total gene counts from this analysis. A total of 6277 genes were detected within the macrophage cluster and tested for differential expression. Of 6277 genes, 929 genes were downregulated, 1101 were upregulated,\ and 4247 genes were unchanged. 26/929 downregulated genes and 23/1101 upregulated genes were PRGs. **d**, Volcano plot displaying the significantly downregulated (sky blue) and upregulated (orange) genes. DE PRGs are highlighted and colored (navy). **e**, DotPlot of all DE PRGs in macrophages displayed as average expression aggregated between conditions (control vs. AAA).

### Metabolic phenotype and phagocytic function of macrophages are altered by LDLR expression in human AAAs

To further elucidate the role of PCSK9 in regulating macrophage function in AAA formation, we performed single-cell pathway analysis (SCPA) to identify cellular functions associated with PCSK9 inhibition.^43,44^ To investigate the effect of PCSK9 inhibition in macrophages, we identified macrophages with high expression of LDLR as a surrogate condition for macrophages after PCSK9 inhibition, since PCSK9 binds LDLR to promote lysosome-mediated degradation (**Fig. 3A** and **Supplementary Table 2**). Using SCPA, we tested curated gene sets of cellular functions (MSigDB C5 GO:Biological Process) between LDLR(+) and LDLR(-) macrophages. Gene sets regulating metabolism and phagocytosis were differentially regulated (**Fig. 3B-C**). Notably, LDLR expression is associated with enrichment of gene sets related to positive regulation of phagocytosis. These findings provide further evidence for a critical role for PCSK9 inhibition in regulating macrophage phenotype during AAA formation, and more importantly, underscores a novel and pivotal mechanistic avenue for PCSK9 inhibitors in attenuating AAA formation by restoring metabolic and inflammatory homeostasis of the aortic wall.

**Fig. 3.**
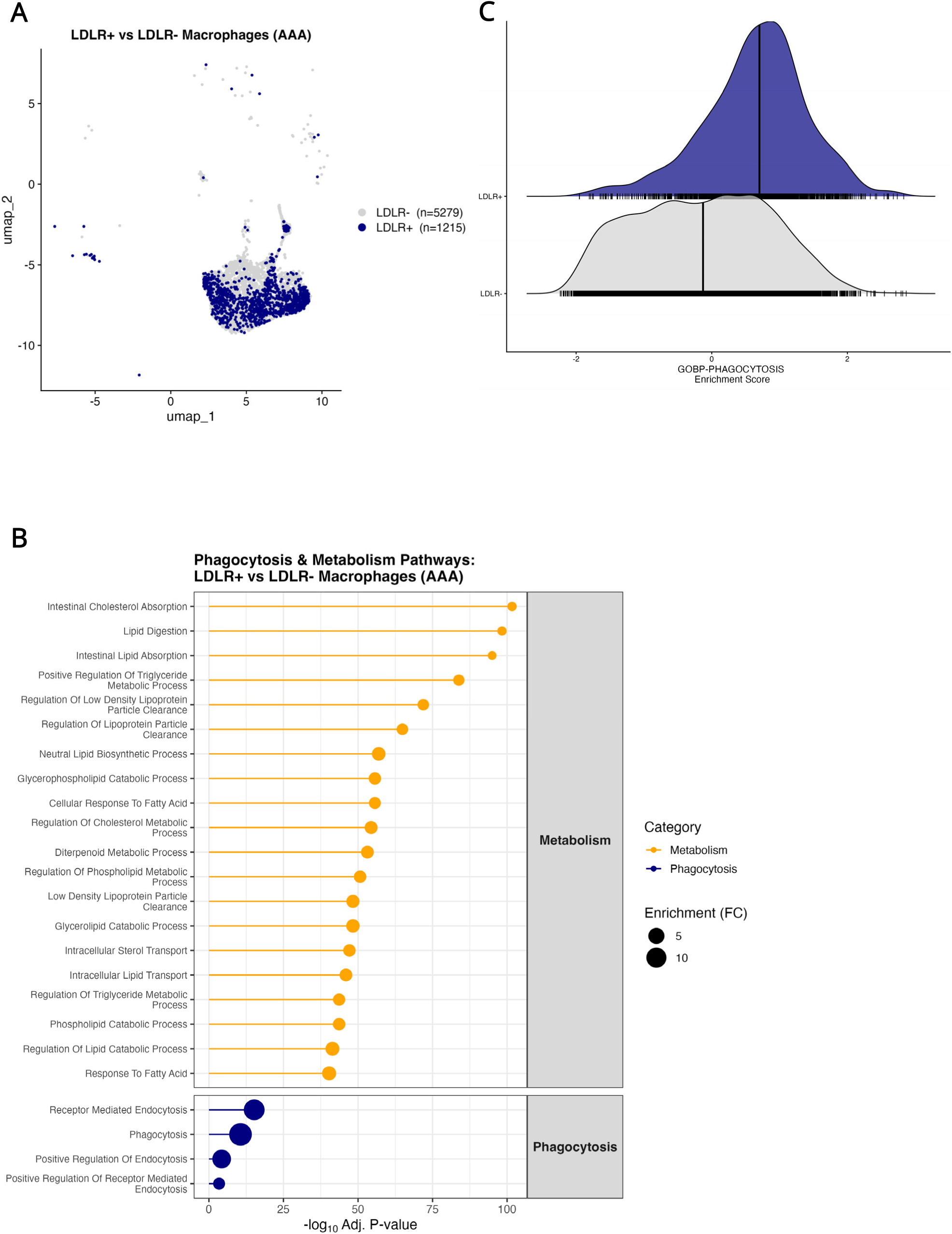
Macrophage LDLR expression modulates metabolic and phagocytic pathways in human AAA. **a**, UMAP projection of macrophages from human AAA samples, classified by LDLR expression (LDLR+, navy, n=1215; LDLR-, grey, n=5279). **b**, Single cell pathway analysis of LDLR+ versus LDLR- macrophages, restricted to GO:BP gene sets. There were 949 differentially expressed gene sets with adjusted p-value <0.01. 106 were related to metabolism. The top 20 differentially expressed metabolic gene sets are displayed, in addition to phagocytosis-related gene sets. Pathways are plotted by statistical significance (-log_10_ adjusted p-value); point size denotes enrichment (fold-change). **c**, Single cell genes set enrichment analysis scores for the GOBP_PHAGOCYTOSIS gene set, shown as ridge density plots by LDLR expression status (LDLR+, navy; LDLR-, grey); vertical lines indicate median values.

### PCSK9 inhibition mitigates aortic inflammation and vascular remodeling in experimental model of AAA formation

Utilizing the established topical elastase AAA model^45^, C57BL/6 (WT) male mice underwent surgeries with or without administration of either vehicle (PBS) or Evolocumab (10 mg/kg) on postoperative days 1 and 7 (**Fig 4A**). A multi-fold increase in mean aortic diameter was observed in elastase-treated mice compared to heat-inactivated elastase-treated (control) mice on day 14 (174.3±8.6% vs. 15.2±5.6%; p<0.0001; **Fig 4B-C**). More importantly, WT mice treated with Evolocumab demonstrated significantly attenuated mean aortic dilation compared to vehicle controls alone (110.94±4.7% vs. 174.3±8.6%; p<0.0001; **Fig 4B-C**).

**Fig. 4.**
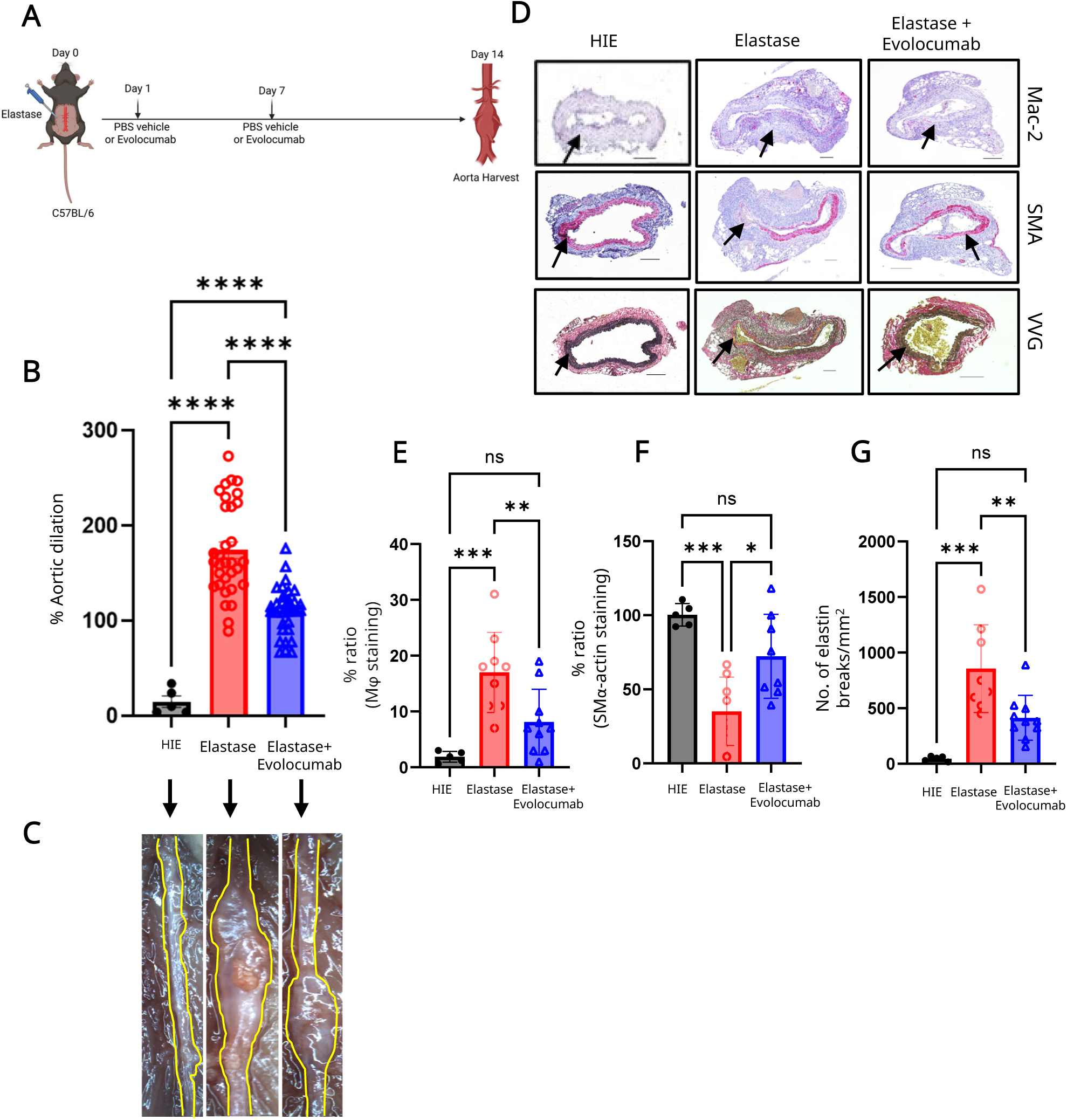
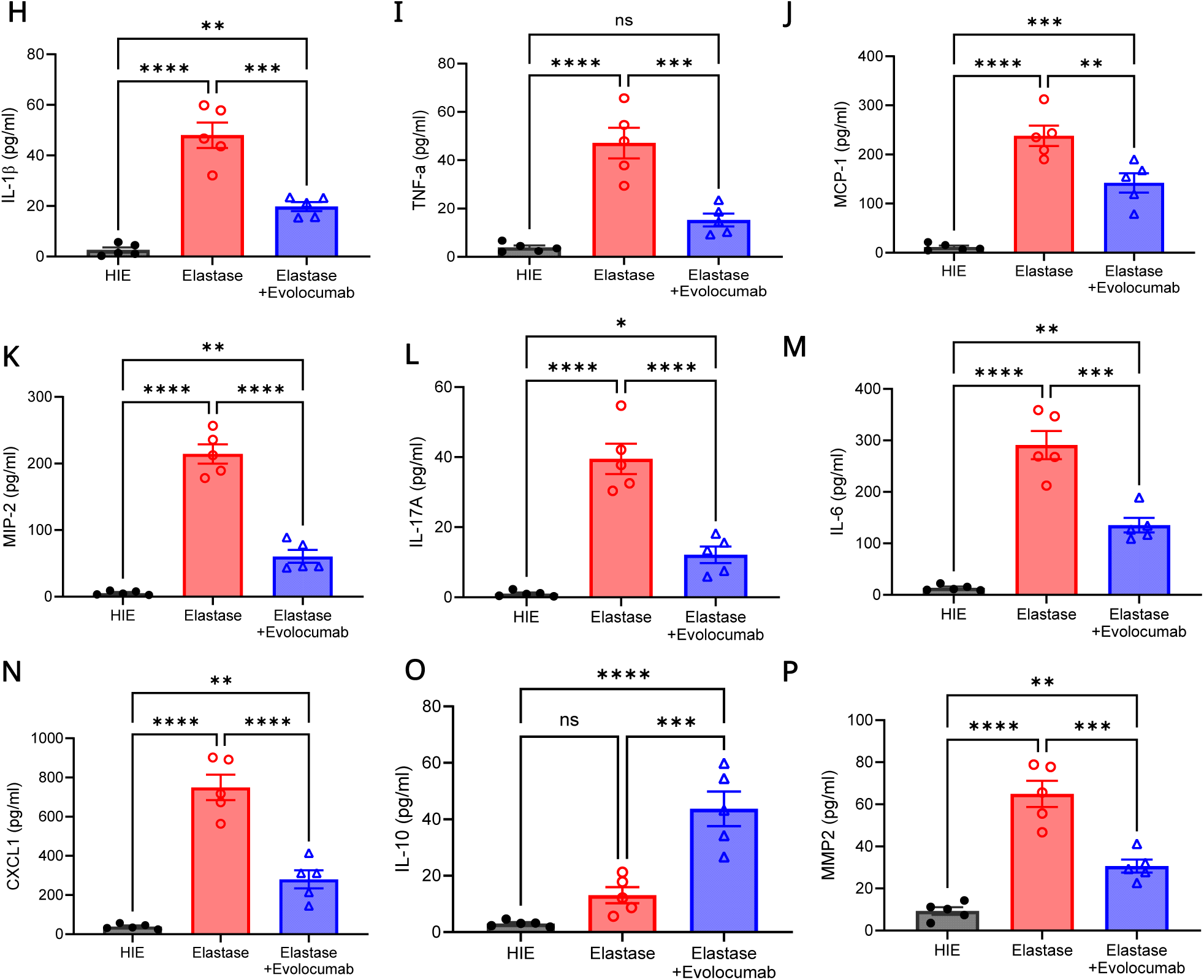
PCSK9 inhibition attenuates aneurysm growth, aortic wall inflammation, and vascular remodeling during AAA formation. **a**, Schematic representation of the murine topical elastase AAA model. **b**, Elastase-treated mice exhibited significant increase in aortic diameter growth compared to heat-inactivated controls, and was significantly attenuated with Evolocumab treatment on day 14. n=5-32/group; ****p<0.0001. **c**, Representative images of aortic diameter measurement. **d-g**, Histologic analysis and quantification of aortic sections demonstrated significantly decreased macrophage infiltration (Mac-2 staining), increased smooth muscle cell α-actin (SM α-actin staining) expression, and decreased elastin fiber disruption (Verhoeff-Van Gieson staining) with Evolocumab treatment. n=5-10/group; ***p<0.001, **p<0.01, *p=0.01, ns=not significant. Scale bar = 100 µm. **h-n**, Pro-inflammatory cytokine expression in aortic tissue was significantly attenuated in the elastase-treated mice after Evolocumab administration compared with vehicle-treated controls. n=5/group; **p<0.005, ***p<0.001, ****p<0.0001. **o**, IL-10 expression was significantly enhanced with Evolocumab treatment compared to untreated controls. n=5/group; ***p<0.001. **p**, MMP2 expression was significantly attenuated in the elastase-treated mice after Evolocumab administration compared to untreated controls. n=5/group; ***p<0.001.

Histological analysis of aortic tissue revealed a significant decrease in macrophage infiltration (8.1±1.9% vs. 17.0±2.4%; p<0.01), increased smooth muscle alpha-actin (SMα-actin) expression (72.4±10.0% vs. 35.1±8.1%; p=0.01), and reduced elastin fragmentation (413.0±64.0 vs. 855.3±139.7; p=0.007 number of breaks/mm^2^) in Evolocumab-treated mice compared to vehicle-treated mice on day 14 (**Fig. 4D-G**). Pro-inflammatory cytokine (IL-1ß, TNF-α, MCP-1, MIP-2, IL-17A, IL-6, and CXCL1) expression in aortic tissue were significantly attenuated, while anti-inflammatory IL-10 expression was significantly enhanced, in elastase-treated mice administered with Evolocumab compared to vehicle-treated controls (**Fig. 4H-O**). Concurrently, matrix metalloproteinase 2 (MMP2) expression was significantly attenuated in Evolocumab-treated mice compared to vehicle-treated mice (**Fig. 4P**).

In separate experiments, mice treated with another PCSK9 inhibitor, Alirocumab^46^, exhibited a significant attenuation in aneurysm growth compared to the elastase treated group (102.6±6.5% vs. 174.3±8.6%; p=0.0002, **Fig. S1A-C**). Additionally, mice treated with recombinant murine PCSK9 (rPCSK9) (0.2µg/µL)^47^ on postoperative days 1 and 7 demonstrated significant enhancement of mean aortic diameter compared to elastase-treated mice (297.5±21.5% vs. 174.3±8.6%; p<0.0001, **Fig. S1A-C**). Collectively, these results demonstrate the ability of PCSK9 signaling to regulate aortic inflammation and vascular remodeling during experimental AAA formation.

### PCSK9 inhibition is associated with pro-efferocytic lipid expression and small molecule metabolites in topical elastase AAA model

Using matrix-assisted laser desorption/ionization (MALDI) mass spectrometry, we investigated identifiable perturbations in the lipidomic and metabolic profile of aortic tissue during AAA formation after Evolocumab treatment. Lipidomic analysis of aortic tissue on day 14 demonstrated increased levels of oxidized and non-oxidized phosphatidylserine (PS(O-45:6) and PS(42:1)) species in the aortic samples of Evolocumab-treated mice compared to untreated controls (**Fig. 5A-B** and **Supplementary Table 3**). Furthermore, phosphatidic acid (PA(42:1), PA(40:1)) and phosphatidylethanolamine (PE(P-40:5), PE(O-40:6)) species were differentially expressed in Evolocumab-treated mice, while lysophospholipid species ( LPI(16:0), LPIP(33:0), LPG(18:1)) were significantly expressed in control AAA tissue compared to the Evolocumab-treated cohort.

**Fig. 5.**
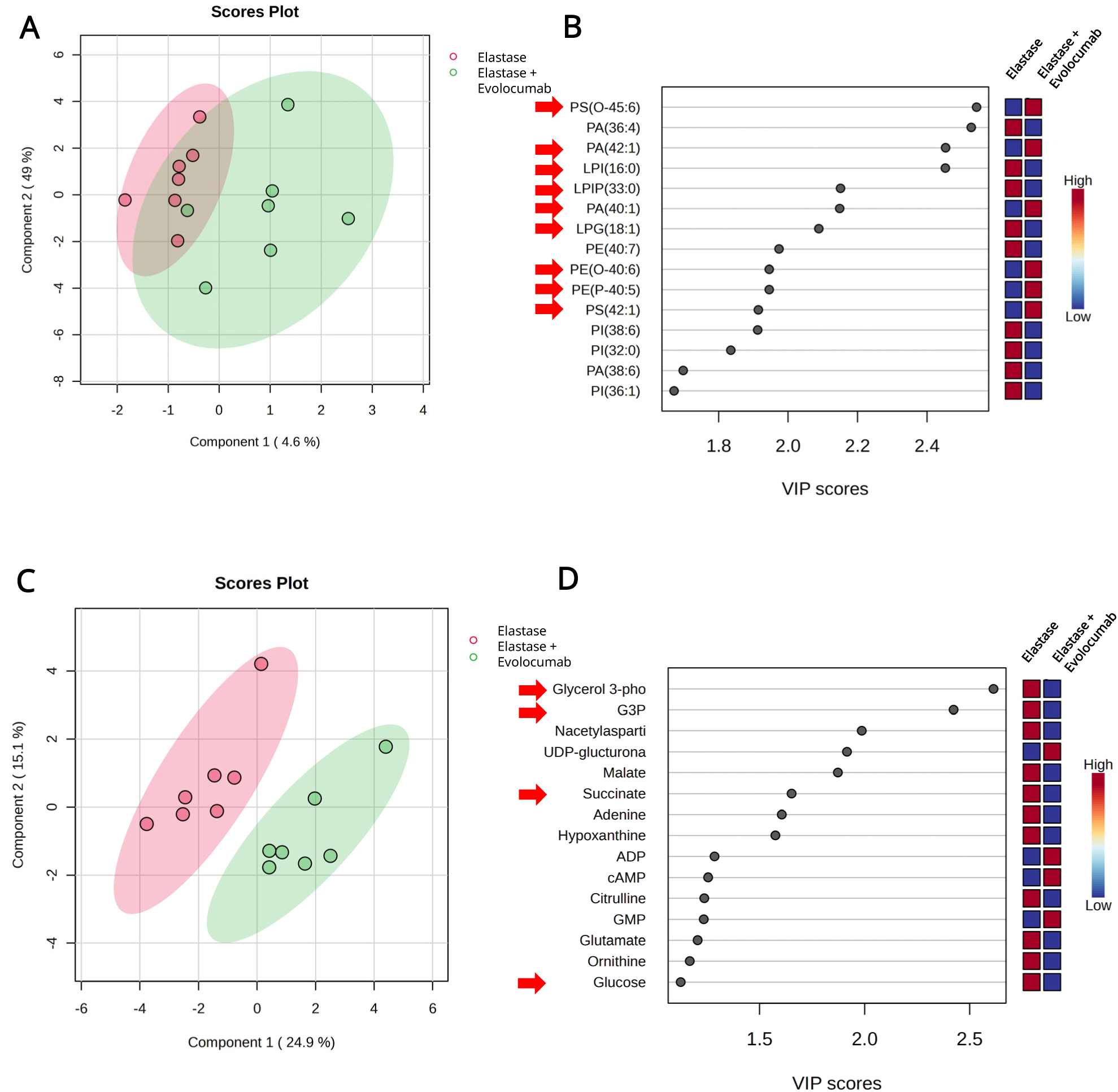
PCSK9 inhibition is associated with pro-efferocytic lipid expression and attenuated anaerobic metabolism. **a**, Untargeted lipidomic score plot from sparse partial least squares discriminant analysis (sPLS-DA), showing separation between vehicle-treated control mice and Evolocumab-treated mice in the topical elastase model on day 14. n=7/group. **b**, Lipids contributing most to group separation ranked by variable importance in projection (VIP) score. Specific referenced lipid species are indicated with red arrows. The accompanying heatmap shows relative metabolite abundance across conditions (red, high; blue, low). PA, phosphatidic acid; PE, phosphatidylethanolamine; LPE, lysophosphatidylethanolamine; PS, phosphatidylserine; PE, phosphatidylethanolamine; LPI, lysophosphatidylinositol; LPG, lysophosphatidylglycerol. **c**, Untargeted small molecule metabolomic score plot from sparse partial least squares discriminant analysis (sPLS-DA), showing separation between vehicle-treated control mice and Evolocumab-treated mice in the topical elastase model on day 14. n=7/group. **d**, Metabolites contributing most to group separation ranked by variable importance in projection (VIP) score. The accompanying heatmap shows relative metabolite abundance across conditions (red, high; blue, low). G3P indicates glyceraldehyde-3-phosphate; ADP, adenosine diphosphate; GMP, guanosine monophosphate; cAMP, cyclic adenosine monophosphate. Specific referenced metabolite species are indicated with red arrows.

Similarly, MALDI analysis identified alterations in the small molecule metabolite profile of aortic tissue in the Evolocumab-treated mice compared to untreated controls (**Fig.5C-D and Supplementary Table 4)**. Glycolytic and anaerobic pathway metabolites and end-products (glyceraldehyde-3-phosphate (G3P), glucose, glycerol-3-phosphate) were differentially expressed in control aortic tissue samples compared to Evolocumab treated mice. Along with shifts toward glycolytic metabolism, select accumulation of key TCA cycle intermediates, such as succinate predominated in control aortic tissues compared to those in Evolocumab treated mice. Collectively, these results suggest a marked change in lipid and glycolytic pathway metabolites during AAA formation which can be altered by Evolocumab treatment.

### PCSK9 inhibition confers protection against preformed AAAs to prevent aortic rupture

Next, we investigated if Evolocumab treatment can confer protection against preformed AAAs and prevent impending aortic rupture, utilizing the advanced stage elastase+ß-aminoproprionitrile (BAPN) murine model of AAA and aortic rupture.^48^ Evolocumab was administered on day 14 and 21 post-surgery, with aortic diameter measurement and harvest performed on day 28 (**Fig. 6A**). Evolocumab treatment resulted in a significant reduction in aortic dilation compared to vehicle-treated controls (182.1±18.7% vs. 347.7±28.2%; p<0.001; **Fig. 6B-C**). Histological analysis and quantification revealed that Evolocumab treatment resulted in a significant decrease in macrophage infiltration (5.2±1.8% vs. 13.1±2.0%; p<0.02), increased smooth muscle alpha-actin (SM α-actin) expression (67.5±4.4% vs. 51.4±2.6%; p<0.02), and reduced elastin fragmentation (541.4±78.1 vs. 918.6±107.8; p<0.02 number of breaks/mm^2^) on day 28 compared to mice treated with elastase+BAPN (**Fig. 6D-G**). Furthermore, Evolocumab treatment significantly mitigated pro-inflammatory cytokine expression (IL-1ß, TNF-α, MCP-1, MIP-2, IL-17A, IL-6, and CXCL1) and increased IL-10 expression in the aortic tissue compared to vehicle-control treated mice (**Fig. 6H-O)**. Moreover, MMP-2 expression was significantly attenuated with Evolocumab treatment compared to untreated controls, suggesting an important role of PCSK9 inhibition on SMC remodeling (**Fig. 6P**).

**Fig. 6.**
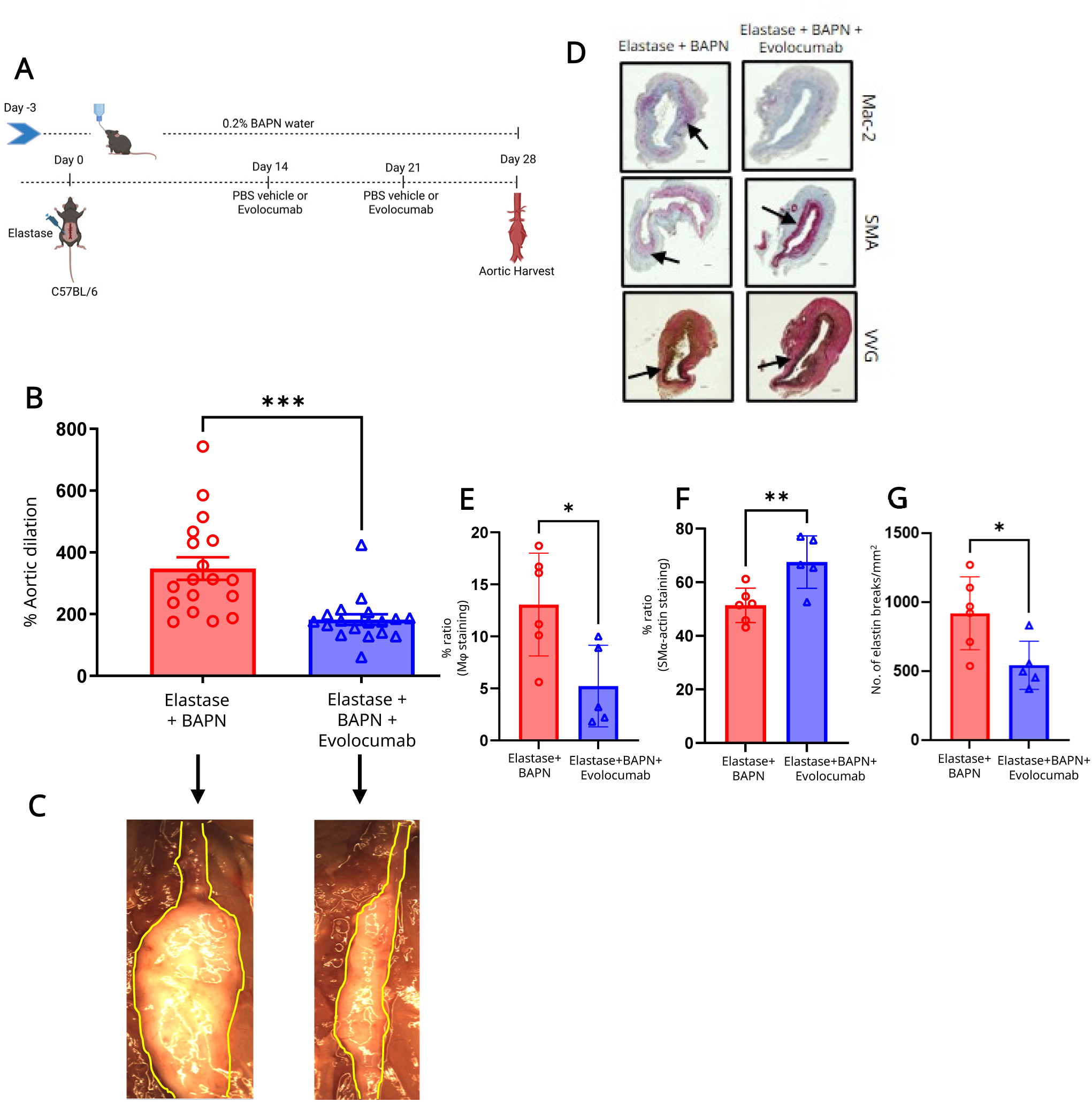
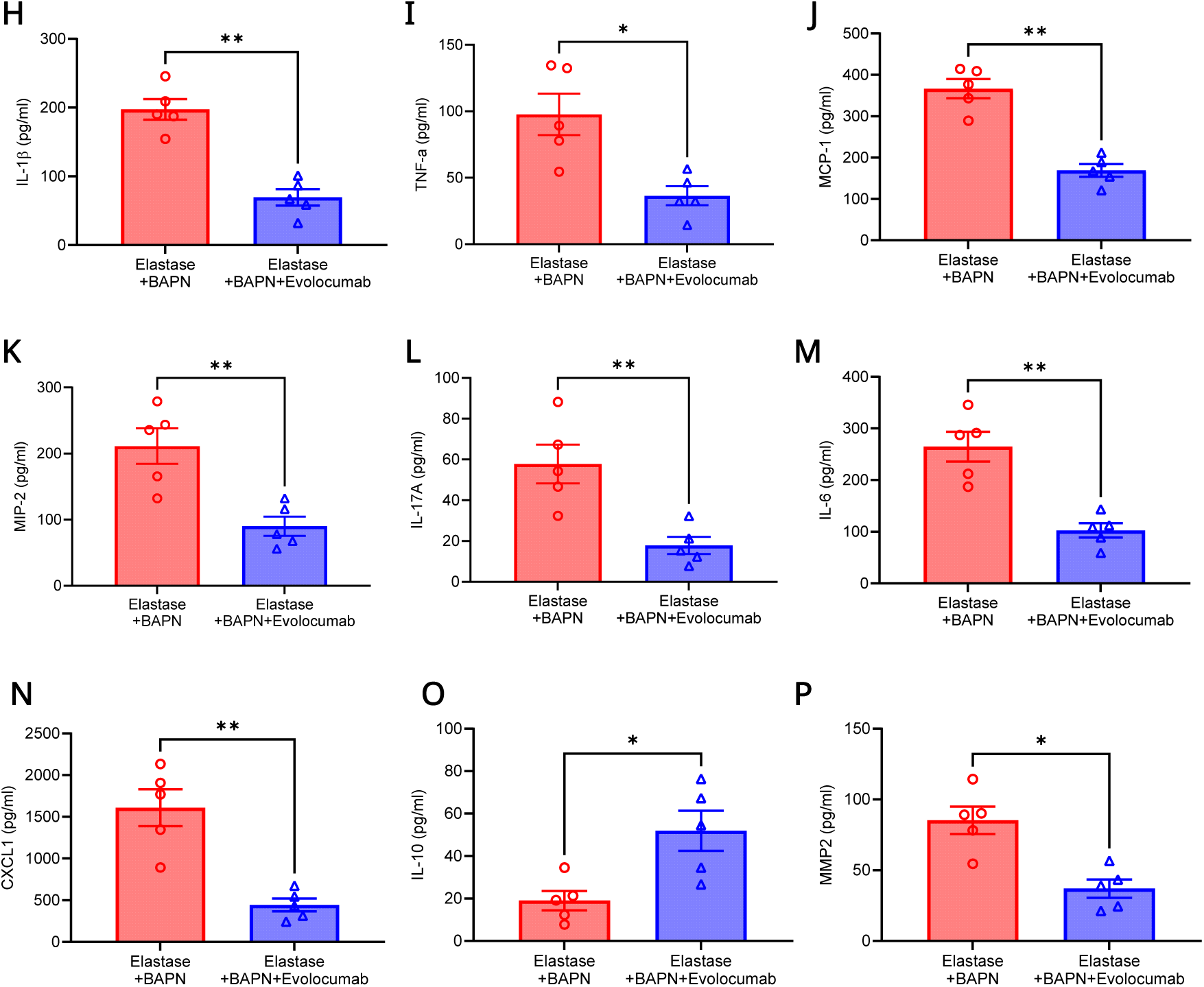
PCSK9 inhibition confers protection against preformed aneurysm in a chronic AAA and aortic rupture model. **a**, Schematic representation of the topical elastase+BAPN (ß-aminopropionitrile) AAA and aortic rupture model. **b**, Administration of Evolocumab significantly attenuated the aortic diameter compared to untreated controls after elastase+BAPN treatment on day 28. n=17-18/group; ***p<0.002. **c**, Representative images of aortic diameter measurement on day 28. **d-g**, Histologic analysis and quantification of aortic sections on day 28 demonstrated significantly decreased macrophage infiltration (Mac-2 staining), increased smooth muscle cell α-actin (SM α-actin staining) expression, and decreased elastin fiber disruption (Verhoeff-Van Gieson staining) after Evolocumab treatment compared to untreated controls. n=5-6/group; *p<0.03, **p<0.01 Scale bar=100 µm. **h-n**, Pro-inflammatory cytokine expression in aortic tissue was significantly attenuated after Evolocumab administration compared to vehicle-treated controls. n=5/group; *p<0.02; **p<0.01 **o**, IL-10 expression was significantly enhanced after Evolocumab treatment compared to untreated controls. n=5/group; *p<0.02. **p**, MMP2 expression was significantly attenuated after Evolocumab treatment compared to untreated controls. n=5/group; *p<0.02.

Moreover, we also used another established experimental angiotensin-II/*ApoE^-/-^* murine AAA rupture model^49^ to confirm the protective effects of PCSK9 inhibition (**Fig. S2A**). The mean aortic diameter was significantly attenuated in mice treated with Evolocumab compared to vehicle-treated control mice on day 28 (87.8±29.3% vs. 256.9±2.3%; p<0.0001; **Fig. S2B-C**). Death from aortic rupture was confirmed via autopsy and mice after Evolocumab treatment experienced a non-significant decrease in rupture-related mortality (80% vs. 61%; log-rank p=0.1; **Fig. S2D**). In summary, our results confirm the ability of PCSK9 inhibition on mitigating pre-formed AAA formation in multiple established murine models by decreasing aortic inflammation and vascular remodeling.

### PCSK9 inhibition enhances MerTK-dependent efferocytosis in macrophages

To elucidate the mechanistic signaling downstream of PCSK9-mediated immunomodulation, we investigated the clearance of apoptotic cells by macrophages through efferocytosis, using both *in vivo* and *in vitro* experiments. Aortic tissue of Evolocumab-treated C57BL/6 mice demonstrated a significantly enhanced clearance of apoptotic cells by macrophages compared to untreated controls on day 7 (32.9±4.0% vs. 21.4±1.9%; p=0.04; **Fig. 7A-C**). The enhanced efferocytosis was accompanied by a significant reduction in apoptotic SMCs in Evolocumab-treated aortic tissue compared to untreated controls (17.6±2.6% vs. 30.8±3.5%; p=0.02; **Fig. 7D-E**). Next, we analyzed if Evolocumab mediated protection is regulated by MerTK-dependent efferocytosis. Evolocumab-treatment attenuated AAA formation in B6:129 mice compared to untreated controls (55.6±14.5% vs. 134.7±12.9%; p<0.04), but failed to attenuate AAA formation in MerTK^-/-^ mice (168.4±27.2% vs. 55.6±14.5%; p<0.001; **Fig. 7F-H**). Additionally, Evolocumab-dependent enhancement of efferocytosis was inhibited in MerTK^-/-^ mice compared to Evolocumab-treated B6:129 controls (3.9±1.5% vs. 7.7±0.7%; p=0.01; **Fig. 7I-K and Fig. S3**). Similarly, Evolocumab-treatment attenuated apoptotic SMCs in B6:129 controls compared to those without treatment (6.8±0.7% vs. 16.8±1.9%; p<0.01), but failed to do so in MerTK^-/-^ mice (28.8±5.1% vs. 6.8±0.7%; p<0.0001; **Fig. 7L-M**).

**Fig. 7.**
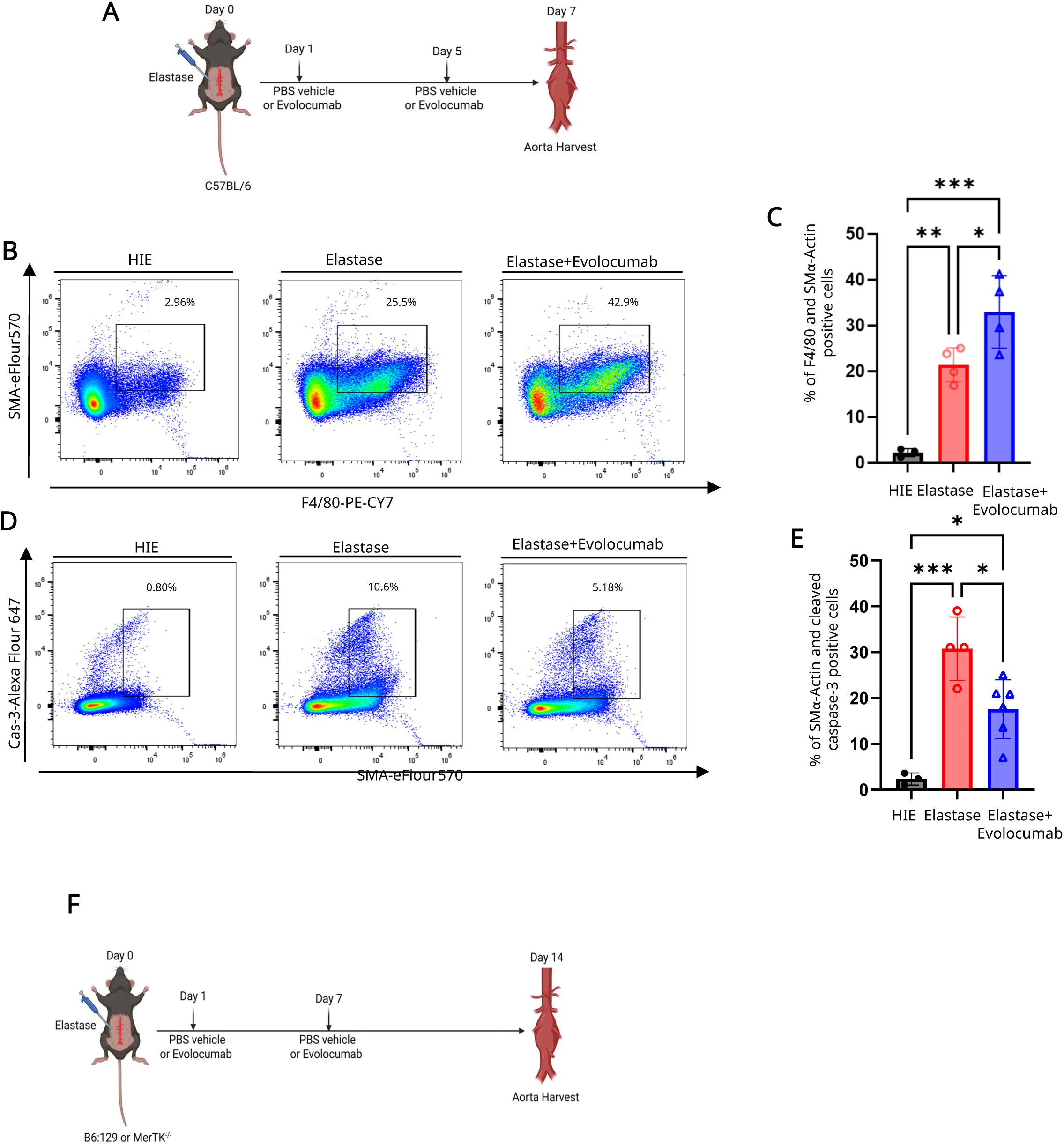
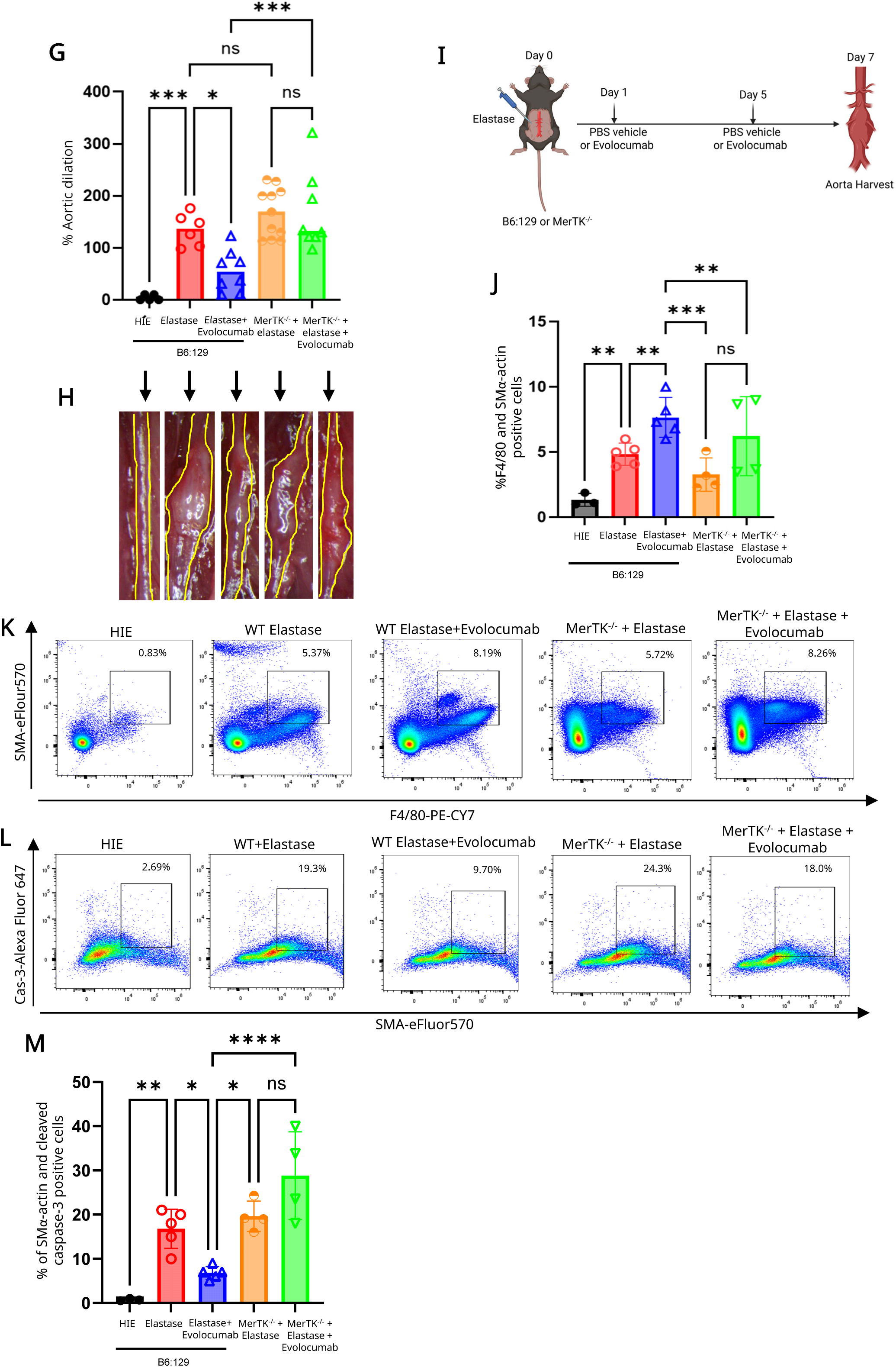
Evolocumab treatment enhances macrophage-dependent efferocytosis *in vivo* during mitigation of AAA formation. **a**, Schematic diagram of the *in vivo* efferocytosis experimental design. **b-c**, Flow cytometry analysis and quantification showed a significant increase in macrophage-dependent efferocytosis in aortic tissue of mice treated with Evolocumab compared to untreated controls. n=4/group. *p<0.04, **p=0.004, ***p<0.001. **d-e**, Flow cytometry analysis and quantification showed a significant decrease in cleaved caspase-3^+^ SMCs after Evolocumab treatment compared to untreated controls. n=4/group; *p<0.04, **p=0.004, ***p<0.001. **f**, Schematic depiction of topical elastase model using MerTK^-/-^ and respective B6:129 controls. **g-h**, Evolocumab administration significantly attenuated increase in aortic diameter in B6:129 mice, but not in MerTK^-/-^ mice. n=5-11/group. *p<0.04, ***p<0.001, ns=not significant. Representative images of respective aortic diameters on day 14 are shown. **i**, Schematic diagram representing *in vivo* efferocytosis assessment on day 7. **j-m**, Flow cytometry analysis and quantification showing increase in macrophage-dependent efferocytosis and decrease in cleaved caspase-3^+^ apoptotic SMCs after Evolocumab treatment in B6:129 mice, but not in MerTK^-/-^ mice treated with Evolocumab. n=3-5/group; *p<0.05, **p<0.01, ***p<0.001 and ****p<0.0001, ns=not significant.

Furthermore, *in vitro* studies were performed using peritoneal macrophages from C57BL/6 mice, with or without pre-treatment with Evolocumab, followed by exposure to apoptotic murine aortic SMCs (MOVAS cells) (**Fig. 8A**). Efferocytosis of MOVAS cells was assessed through quantifying the percentage of macrophages (F4/80^+^) that phagocytosed PKH67-labeled MOVAS cells (**Fig. 8B-C**). Evolocumab treatment significantly enhanced the efferocytic function of macrophages compared to vehicle-treated controls (43.6±2.1% vs. 37.0±0.7%; p=0.04), which was inhibited in the presence of MerTK inhibitor (27.4±0.5% vs. 43.6±2.1%; p<0.0001; **Fig. 8B-C and Fig. S4A-B**). Conversely, recombinant PCSK9-treatment significantly decreased macrophage efferocytic function compared to Evolocumab-treatment (27.7±0.6% vs. 43.6±2.1%; p<0.0001). Moreover, *in vitro* quantification of Arginase1 (ARG1) activity was performed to ascertain the degree of M2 macrophage polarization occurring across treatment conditions (**Fig. 8D**). Macrophages treated with Evolocumab had significantly higher ARG1 concentrations compared to untreated controls (61.0±2.48 vs. 49.13±0.97 ng/mL; p<0.02). Additionally, the Evolocumab dependent increase in ARG1 expression was mitigated with MerTK inhibition (30.75±4.13 vs. 61.0±2.48; p<0.0001). Furthermore, treatment with recombinant PCSK9 protein significantly attenuated ARG1 expression (45.63±0.9 vs. 61.0±2.4 ng/ml; p<0.01). Collectively, these results demonstrate that PCSK9 inhibition regulates macrophage-specific efferocytosis of apoptotic SMCs via MerTK receptors and enhances M2 polarization, restoring metabolic homeostasis of the aortic wall and mitigating AAA formation (**Fig. 8E**).

**Fig. 8.**
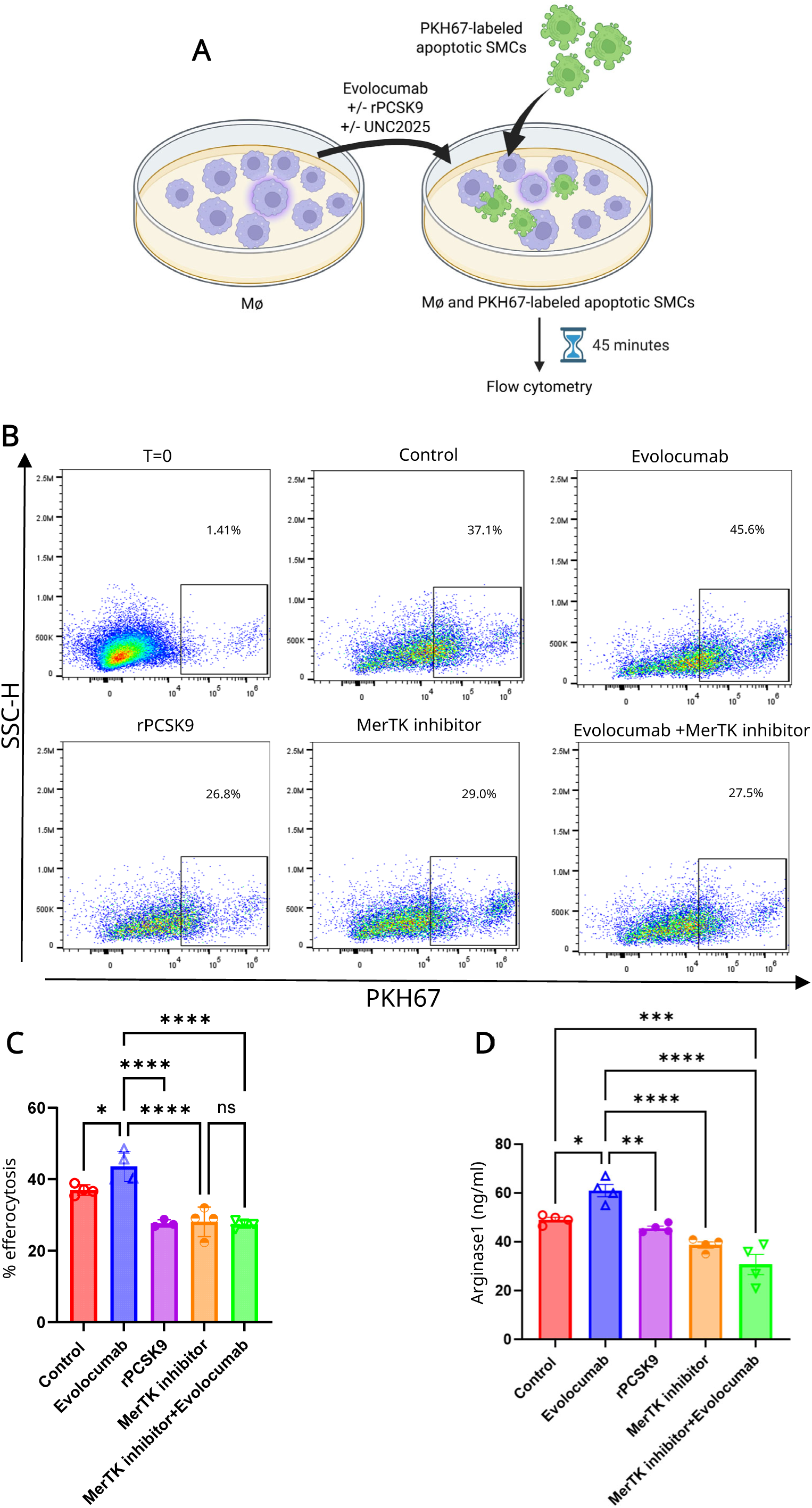
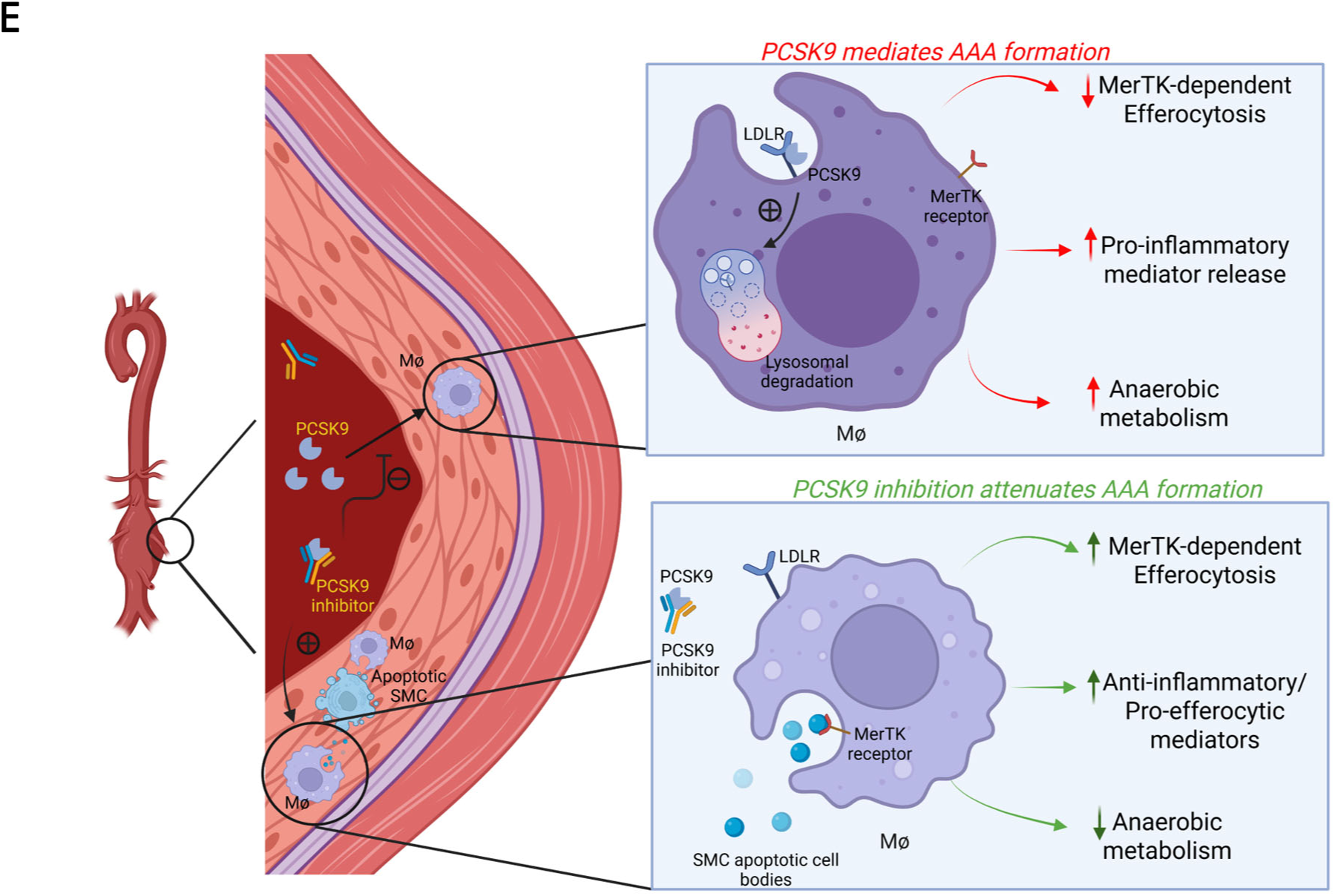
Evolocumab enhances macrophage-dependent efferocytosis *in vitro* in a MerTK-dependent mechanism. **a**, Schematic representation of *in vitro* efferocytosis experimental design. **b-c**, Evolocumab treatment significantly enhances macrophage-mediated efferocytosis compared to vehicle-treated controls in a MerTK-dependent manner. Treatment with recombinant PCSK9 significantly mitigates macrophage-dependent efferocytosis. n=3-4/group; *p<0.04, **p<0.006, ****p<0.0001, ns=not significant. **d,** Evolocumab treatment significantly enhances ARG1 expression in macrophages whereas recombinant PCSK9 significantly decreases ARG1 expression. n=4/group; *p<0.02, **p<0.01, ***p<0.001, ****p<0.0001. **e**, Schematic diagram representing PCSK9 mediated effects in AAA formation, including decreased MerTK-dependent efferocytosis, increased pro-inflammatory mediator release (cytokines, lysophospholipid species, TCA cycle intermediates), and increased anaerobic metabolism (glyceraldehyde-3-phosphate, glucose, glycerol-3-phosphate) within the aortic wall. Conversely, PCSK9 inhibition attenuates AAA formation through enhanced MerTK-dependent efferocytosis, increased anti-inflammatory mediator release (IL-10, phosphatidylserine, phosphatidic acid, and phosphatidylethanolamine species), and diminished anaerobic metabolism.

## Discussion

This study demonstrates that PCSK9 signaling promotes AAA formation by driving aortic inflammation and vascular remodeling observed during AAA pathogenesis. Conversely, clinical grade inhibition of PCSK9 using two different pharmacological medications (Evolocumab and Alirocumab), results in attenuation of aortic inflammation and AAA growth by enhancing macrophage-dependent efferocytosis of apoptotic SMCs and restoring metabolic homeostasis within the aortic wall. The clinical relevance of these findings is reinforced by a correlative retrospective study showing a significant association between decreased mortality and rates of AAA growth in patients on PCSK9 inhibitor therapy. Second, single-cell RNA-sequencing analysis of human AAA tissue demonstrated dysregulation of several PCSK9-related genes within macrophages, specifically involved with inflammatory modulation and phagocytic function within AAA. Third, complimentary single-cell pathway analysis from a human gene set database also revealed dysregulations in macrophage metabolism, inflammatory regulation, and phagocytic function in response to variable expression of LDLR, further supporting the direct impact of PCSK9-mediated degradation of LDLR on reprogramming macrophage phenotype in AAA.

Pharmacologic treatment with PCSK9 inhibitors independently mitigated AAA growth across multiple established preclinical murine models of AAA^45,48–51^, and consistently showed sustained reductions in degenerative vascular remodeling, macrophage infiltration, and pro-inflammatory cytokine expression. Finally, *in vivo* and *in vitro* experiments demonstrated the previously unrecognized ability of PCSK9 inhibition to upregulate macrophage-dependent efferocytosis of apoptotic SMCs in a MerTK-dependent manner, and enhance M2 macrophage polarization, resulting in attenuated aortic inflammation and vascular remodeling during AAA formation.

Perturbations in lipid biology have been intricately associated with dysregulation of vascular pathologies, such as atherosclerosis and AAA formation.^52–54^ Recent studies have postulated that dysregulated lipid metabolism, including lipid peroxidation, is associated with cell death processes, such as apoptosis and ferroptosis of aortic SMCs, leading to aortic wall remodeling and AAA development. ^15,16,55^ Lipidomic analysis of aortic tissue from Evolocumab-treated mice have revealed significantly elevated levels of oxidized and non-oxidized phosphatidylserine species. Externalization of phosphatidylserine is a known hallmark of apoptosis but oxidized phosphatidylserine variants have been associated with more efficient efferocytosis with limited inflammation through modulation of macrophage-specific inducible NO synthase (iNOS) and IL-1ß transcription through LPS-mediated signaling.^56–58^ Additionally, we observed that phosphatidic acid and phosphatidylethanolamine species were preferentially expressed in Evolocumab-treated aortic tissue. These have been previously described as intermediary byproducts of palmitoylethanolamide and oleoyethanolamide synthesis, which signal to enhance M2 macrophage polarization and subsequent apoptotic cell body ingestion.^59^ Furthermore, lysophospholipid species were increasingly prevalent in AAA tissue compared to Evolocumab-treated cohorts. These abnormal lipid metabolites function as damage-associated molecular patterns (DAMPs), which further augment the inflammatory milieu impacting the aortic wall in AAA and impairing macrophage function.^60^

Metabolomic profile analysis comparing aortic tissue of Evolocumab-treated cohorts revealed perturbations reflective of enhanced macrophage polarization and efferocytosis secondary to PCSK9 inhibition. Pro-resolving M2 macrophages are more metabolically efficient for efferocytosis than pro-inflammatory M1 macrophages, utilizing apoptotic body-derived substrates preferentially for oxidative phosphorylation.^61,62^ Our data shows that glycolytic and anerobic pathway intermediates were elevated in control aortic tissue, reflecting shifts toward less-efficient means of energy acquisition. Additionally, TCA cycle derangements occur with M1 macrophage polarization, resulting in accumulation of key intermediates, such as succinate, which furthermore function in intracellular and extracellular signaling pathways to propagate inflammation.^63–65^ The enhanced expression of these metabolites predominated in control AAA tissues compared to aortic tissues from mice treated with Evolocumab, signifying alterations in the metabolic environment of the aortic wall, concordant with improved macrophage-dependent efferocytosis observed within the previously described *in vivo* and *in vitro* experiments.

These findings emphasize the intricate balance between homeostasis and immunometabolism in the lipid and metabolite profile impacts immune cell and resident cell phenotype. The ability of PCSK9 inhibitors to immunomodulate these metabolic alterations to synergistically mitigate chronic inflammation and degenerative vascular remodeling offers a unique perspective in the context of vascular diseases. Moreover, previous experimentation has supported the role of endogenous PCSK9 to promote macrophage migration and pro-inflammatory mediator release in aneurysm aortic walls, as well as different spectra of vascular disease, including atherosclerosis, atherosclerotic necrotic core formation, and aortic dissection.^11,23, 66–70,71^ However, the dysregulation of efferocytosis to clear dead cell debris in the aortic milieu can propagate thrombus formation and chronic inflammation, and remains to be delineated specifically in the context of PCSK9 inhibition.

In diverse disease processes, PCSK9 has been shown to promote macrophage phenotypic polarization toward an activated, pro-inflammatory M1 state.^72,73^ Compared to their M2 counterparts, M1 macrophages have diminished metabolic and efferocytic efficiency. MerTK is a macrophage cell-surface receptor that is a key regulator of macrophage-dependent efferocytosis, through its function to effectively recognize and bind preserved moieties (i.e., phosphatidylserine) associated with apoptotic cell bodies via circulating bridge molecules (Gas6, ProteinS, Tubby, and Tubby-like protein).^74,75^ Previous studies have identified associations with knock down or knockout MerTK function with deleterious phenotypic changes in vascular disease, including increased atherosclerotic necrotic core formation, SMC apoptosis, and deficient macrophage-dependent efferocytosis.^76,77^ Furthermore, increased M2 macrophage expression of MerTK is associated with attenuated atherosclerotic disease burden in patients with coronary artery disease.^78^ These results are the first to describe the role of MerTK receptors on macrophages to regulate efferocytosis of apoptotic SMCs in the context of AAAs and demonstrate the ability of Evolocumab to significantly enhance macrophage-dependent efferocytosis. Additionally, enhancement of M2 macrophage polarization mediated through Evolocumab was observed via increased ARG1 concentrations in macrophages. Historically, ARG1 expression has been viewed as a predominant marker of M2 macrophage phenotype, and is associated with anti-inflammatory and vascular remodeling functions.^79,80^ This previously unrecognized paradigm of PCSK9-mediated, lipid-independent pathways via MerTK receptor mediated efferocytosis by macrophages represents a significant advancement in our understanding of inflammation and dysregulated resolution of chronic vascular pathologies and is likely to have relevant implications in other diseases apart from AAAs.

There are several limitations to consider in this study. First, lipidomic and metabolomic analyses were performed on murine aortic tissue samples coinciding with significant AAA formation. However, the sequential alterations in lipid and metabolite expression are not completely captured to delineate the full spectrum of metabolic perturbations that occur throughout aneurysm pathogenesis, as well as the relative alterations conferred by PCSK9 inhibition. Furthermore, experimental studies were not performed in the setting of hypercholesterolemia, which is clinically common in the setting of AAA. Although previous studies using PCSK9-AAVs^81,82^ have shown the role of hyperlipidemia in the context of AAAs, our goal was to delineate lipid-independent pathways that could determine macrophage reprogramming secondary to PCSK9-mediated signaling. Also, it is possible that dysregulations of PCSK9-related genes or Ldlr-related pathways in other cell types, such as SMCs and endothelial cells, may also significantly contribute to the pathogenesis of AAA. Additionally, the retrospective clinical dataset represents a single-center study with associated limitations with respect to long-term follow-up, which likely influenced our ability to ascertain significant differences in mortality impact. However, the marked effect on mitigating AAA growth and matched mortality rates are promising to support the feasibility of future prospective studies with PCSK9 inhibitors in AAA patients.

In summary, these results emphasize the importance of the pharmacologic inhibition of PCSK9 to enhance macrophage-dependent efferocytosis of apoptotic SMCs and contribute to the inhibition of aortic inflammation and remodeling during AAA formation. Our results highlight the multimodal impacts of PCSK9 inhibition on AAA attenuation, through both lipid and LDLR-dependent and independent mechanisms, with translated findings in adjunctive retrospective clinical and preclinical cohorts. These findings support the utility of a multi-center prospective clinical trial to investigate the repurposing of PCSK9 inhibitor therapy as a viable therapeutic modality for patients with AAA.

## Acknowledgements

We thank Tabitha Randi for technical assistance and laboratory maintenance in the Aortic Aneurysm research laboratory at University of Florida. We also thank Roberto Ribas, Reece C. Larson, Alison M. Ryan, Charles Soto, and Franco Bucco Paolasso at CASBR (Center for Advanced Spatial Biomolecule Research) at University of Florida. Additionally, we also thank the Society for Vascular Surgery (SVS) and the family of Dr. James S.T. Yao for presentation of these findings at the 2026 SVS Vascular Annual Meeting in Boston, MA.

## Supplementary Material

Supplementary material (Figures S1-S4 and Tables S1-S4) are available online.

## Funding

This work was supported by the following National Institutes of Health grants NIH R01 HL138931 and RO1 HL153341 (GRU and AKS), NIGMS postgraduate training grant T32 HL160491 (GRU), NIH R01AG066653, R01CA266004, R01AG078702, R01CA288696, RM1NS133593 (RCS), and the Frederick A. Coller Surgical Society (MF).

## Data Availability

All relevant data are included in the manuscript or as Supplementary material.

## Conflict of Interest

The authors declare no conflicts of interest in connection with this article.

## Materials and Methods

### Human clinical data analysis

For clinical data analysis, aortic aneurysm patients were identified utilizing ICD-10 codes (I71.40, I71.43, I71.4, I71.41, I71.42) for AAA between 2015-2023 from the University of Florida Integrated Data Repository (IRB202301533). Comorbidity and lipid therapy status (no therapy, statin therapy alone, PCSK9 inhibitor therapy alone) were extracted and manually confirmed (MF, GS, WU, GG). Patients with an incomparable number of computed tomography scans (<2) were excluded from analysis. Patients with documented connective tissue disorders, aortic dissections, or those that underwent surgical repair of their aneurysm between imaging intervals were also excluded. Manual review and measurement of AAA size in both transverse and anteroposterior diameter were performed by one standardized reviewer (MF). Patients with maximum aortic diameter growth exceeding >2 mm between scans were designated as experiencing AAA growth; while maximum diameter changes ranging from −2 to 2 mm were defined as AAA stabilization. AAA regression was defined as a decrease in maximum aortic diameter exceeding 2 mm between scan intervals. Nearest-neighbor propensity score matching was performed to correct for comorbidity burden between treatment groups, using hypertension, diabetes, smoking status, and atherosclerosis (defined through ICD10 codes: I70.0-I70.9) as covariates, with a primary endpoint of overall mortality. Raw and matched comparisons were performed using Kruskal-Wallis (continuous variables) and Fisher’s exact tests (categorical variables). Kaplan-Meier estimates were used to determine long-term survival estimates based on therapy. Continuous variables are presented as median [IQR] (range); categorical variables presented as N(col%). A p<0.05 was considered significant.

### Human single cell RNA sequencing analysis

Publicly available single-cell RNA-sequencing datasets of human abdominal aortic aneurysms (AAA; n = 7) and healthy control aortas (n = 5) were obtained from the Gene Expression Omnibus (GSE226492, GSE152583)^31,32^. Raw count matrices were processed in Seurat (v5.4.0)^83^ with standard quality-control filtering to exclude low-quality cells and likely doublets, and datasets were integrated using Harmony^84^ to correct for batch effects. Dimensionality reduction was performed by principal component analysis and uniform manifold approximation and projection (UMAP), and cell clusters were identified using the Louvain algorithm and annotated by canonical lineage markers. The macrophage population was isolated for all subsequent analyses. Differential gene expression between AAA and control macrophages was assessed using the FindMarkers function (Wilcoxon rank-sum test), restricted to genes detected in at least 10% of cells in either group, with no fold-change pre-filter applied. Genes were considered significant at an adjusted p-value < 0.05 and absolute log₂ fold change > 0.25. To identify PCSK9-related genes (PRGs), a curated gene list was obtained from GeneCards^85^ (relevance score ≥ 1) and intersected with the differential expression results. To characterize the biological functions associated with LDLR expression, AAA-derived macrophages were classified as LDLR-positive or LDLR-negative based on detectable LDLR transcript expression. Differences in gene set enrichment between LDLR-positive and LDLR-negative macrophages were evaluated using Single-Cell Pathway Analysis (SCPA) with gene sets from the MSigDB C5 GO:Biological Process collection (species = human).^43,44^ Differentially enriched gene sets were defined at an adjusted p-value < 0.01. Single-sample gene set enrichment analysis (ssGSEA), implemented in the escape package, was used to compute per-cell enrichment scores for phagocytosis-related gene sets and to compare their distributions between LDLR-positive and LDLR-negative macrophages.

### Murine AAA models

Male wild-type C57BL/6 mice (8-12 weeks) were obtained from Jackson Laboratory (Bar Harbor, ME). Additionally, male wild-type B6:129 and B6:129-Mertk^tm1Grl/J^ (global MerTK^-/-^) mice were obtained from Jackson Laboratory (Bar Harbor, ME). The animals were housed under controlled conditions, including: 70°F temperature, 50% humidity, and a 12-hour light–dark cycle, in accordance with institutional animal care guidelines. Mice had access to a standard chow diet and drinking water. All experimental procedures were approved and conducted in accordance with the Institutional Animal Care and Use Committee at the University of Florida (protocol #201910902). Two murine elastase-treatment models of AAA formation were used, as previously described.^46,48,50,51^ Mice were anesthetized with inhaled isoflurane (1-2% mg/mL at 0.5 mL/min) and 3.25 mg/kg subcutaneous Ethiqa XR (Fidelis Animal Health, North Brunswick, NJ) and 1-2 mg/kg 0.5% bupivacaine (Hospira Inc., Lake Forest, IL). The abdominal aorta was exposed through a midline laparotomy, and the infrarenal aorta was then circumferentially dissected and topically treated for 3 minutes with 5 µL of elastase (0.4 U/mL, type I porcine pancreatic elastase; Sigma-Aldrich, St. Louis, MO), or heat-inactivated elastase (HIE), which functioned as a control.

In the advanced stage AAA model, mice were pre-treated with supplementation of 0.2% ß-aminopropionitrile (BAPN) (Sigma-Aldrich, St. Louis, MO) administered within drinking water, starting 72 hours prior to aneurysm induction with topical elastase through day 28 post-surgery. Treated aortic segments were harvested either on day 14 (topical elastase model) or day 28 (advanced stage/rupture model), and snap-frozen in liquid nitrogen for molecular analysis or preserved in paraformaldehyde for immunohistochemistry. Aortic diameter was assessed through video micrometry (AmScope, Irvine, CA), and the degree of aortic dilation was determined through using the formula: [(maximal aortic diameter – baseline diameter)/baseline diameter] x 100. Aortic dilation ≥100% indicated AAA formation.

We also used the angiotensin-II model of AAA rupture for this study, as previously described.^45,50,51^ Briefly, male *ApoE*^-/-^ mice ranging 8-12 weeks of age were pre-treated for 4-weeks duration with a high-fat diet (D12492i, Research Diets Inc., New Brunswick, NJ) prior to subcutaneous implantation of osmotic pumps (Alzet 2004; Durect, Cupertino, CA) containing angiotensin II (Ang II, 2000 ng/kg/min, Sigma-Aldrich, St. Louis, MO). The high-fat diet was continued throughout the study duration. On day 28, aortic tissue was removed in its entirety, from the aortic root en-bloc with the heart, to the level of the iliac bifurcation, and aortic diameter measurements were obtained. Again, aortic dilation ≥100% was indicative of aneurysm formation.

### Histology

Following harvest, resected aortic segments were fixed overnight in 10% zinc-buffered formalin, and subsequently dehydrated through a graded ethanol series, before embedding in paraffin. Tissue sections were stained for immunohistochemical analysis using the following primary antibodies: anti-mouse Mac-2 for macrophages (1:5000; Cedarlane Laboratories, Burlington, ON, Canada; catalog no. CL8942AP), anti-mouse α-smooth muscle actin (α-SM, 1:000; Sigma, St. Louis, MO; catalog no. A5691), and Verhoeff-Van Gieson (VVG) for elastin (Polysciences, Inc., Warrington, PA; catalog no. 25089-1). Quantification of elastin degradation was determined through counting the number of elastin fiber breaks per mm^2^ using Fiji (ImageJ, NIH), counts were averaged and depicted graphically. Histologic evaluation was conducted on three individual aortic sections per mouse, and quantification was conducted independently via blinded observers. For consistency and accuracy, representative images from multiple animals per group were chosen. Imaging was conducted at 20x magnification with a digital Nikon microscope, equipped with NIS-Elements BR software. Quantification of histology staining intensity was conducted using QuPath version 0.5 (QuPath; University of Edinburgh, Edinburgh, UK) through quantifying the percentage of positively stained areas in relation to the total aortic cross-sectional area.

### Cytokine multiplex assay

Cytokine expression in homogenized murine aortic tissue was quantified utilizing the Bio-Plex Bead Array system with a multiplex cytokine panel assay (Bio-Rad Laboratories, Hercules, CA). Harvested aortic tissues were first snap-frozen in liquid nitrogen. Frozen samples were then ground using a mortar and pestle and homogenized within tissue extraction buffer. The BCA assay was utilized to quantify protein concentration. Equivalent protein concentrations were used for cytokine measurements within the tissue samples.

### Lipidomic and metabolomic analysis

Matrix-assisted laser desorption/ionization (MALDI) mass spectrometry^86^ was utilized to study the lipidomic and metabolomic profile changes to the aortic wall following treatment with Evolocumab. Aortic tissue was harvested and embedded in frozen deionized water. Fresh frozen tissue was then cut onto a Fisherbrand^TM^ Superfrost^TM^ Plus (Fisher Scientific, Pittsburgh, PA) microscope slide at −20°C on a Leica CM1860 cryostat (Leica Biosystems, Buffalo Grove, IL). Tissue sections were then dehydrated in a desiccator for 60 minutes to prevent further metabolism. Slides were then sprayed utilizing a HTX M5 Sprayer (HTX Imaging, Chapel Hill, NC) with N-(1-naphthyl) ethylenediamine dihydrochloride in 70% methanol at 7 mg/mL. Slides were then vacuum dried in a desiccator for 15 minutes. Slides were analyzed by a Bruker timsTOF Flex (Bruker Nano Analytics, Berlin, Germany) with a raster and spot size of 50 µm to image for lipids and small molecules. Laser power was adjusted to 10,000 Hz, 396 laser shots per pixel, and 70% laser energy. Mass acquisition was set from 50-2,000 m/z in negative mode. Following scanning, all slides were imported into Bruker SCiLS lab (Bruker Nano Analytics, Berlin, Germany) for putative peak annotation and MetaboAnalyst 6.0 (MetaboAnalyst, Alberta, Canada) for downstream analysis.

### *In vivo* efferocytosis assay

*In-vivo* efferocytosis assay was performed using C57BL/6, B6:129, and global MerTK^-/-^ that underwent topical elastase AAA induction, followed by intraperitoneal administration of either PBS vehicle or 10 mg/kg Evolocumab on day 1 and 5. On day 7, the aneurysmal infrarenal aorta was excised under sterile conditions and dissected free of periadventitial tissue. The aortic tissue samples were then fragmented into ∼1 mm^3^ sections and digested in type I collagenase (2.5 mg/mL, Worthington Biochemical, Lakewood, NJ) and porcine pancreatic elastase (0.5 mg/mL, Sigma-Aldrich, St. Louis, MO) prepared in DMEM plain medium at 37°C for 40 min and carefully agitated. FBS (10% v/v) was added to tissue digest to arrest enzyme activity, then passed through a 70-µm strainer and placed into cold PBS buffer. Cell suspensions were centrifuged at 400 x g for 5 min at 4°C and washed prior to Live/Dead blue fixable staining (Thermo Fisher Scientific, Waltham, MA) as per manufacturer’s recommendations. Following the Live/Dead blue fixable staining, the cells were further stained in cell staining buffer containing 2% FBS and 2mM EDTA in PBS. Cells were then incubated with anti-mouse CD16/32 (10 µg/mL, Fc Block, BioLegend, San Diego, CA) at 4°C for 10 minutes and then stained at 4°C for 30 minutes with anti-mouse CD45-BV785 (BioLegend, San Diego, CA) and anti-mouse F4/80-PE-Cy7 (eBioscience, San Diego, CA). Following surface antigen staining, cells were fixed in 4% paraformaldehyde for 20 minutes at room temperature. Cells were then washed and permeabilized using intracellular staining buffer and incubated with anti-α-smooth muscle actin (α-SMA)-eFluor 570 (Thermo

Fisher Scientific, Waltham, MA) and anti-cleaved caspase-3-Alexa Fluor 647 (Cell Signaling Technology, Danvers, MA) for 30 min at 4°C in the dark. After staining, cells were washed twice with permeabilization buffer and resuspended in staining buffer prior to data acquisition. Samples were acquired on a Cytek Aurora spectral flow cytometer (Cytek Biosciences, Fremont, CA) and data were analyzed using FlowJo software (BD Biosciences, Franklin Lakes, NJ)

### *In vitro* efferocytosis and Arginase1 assay

*In-vitro* efferocytosis assay was performed using thioglycolate-elicited mouse peritoneal macrophages and apoptotic MOVAS (mouse aortic smooth muscle cells; ATCC, Manassas, VA) cell targets. C57BL/6 male mice were administered 3% Brewer thioglycolate broth (Sigma-Aldrich, St. Louis, MO) intraperitoneally. Peritoneal exudate cells were then sampled through lavage with ice-cold sterile PBS at 72 hours. Harvested cells were placed in RPMI-1640 medium supplemented with 10% fetal bovine serum (FBS) for culture along with 1% penicillin-streptomycin and incubated at 5% CO_2_ atmosphere at 37°C for 2.5 hours. Macrophages were pre-treated with or without Evolocumab (5 µg/mL) for 2 hours prior to the assay. Control groups were also treated with either recombinant PCSK9 (1 µg/mL) (Abcam, Trumpington, UK), or selective inhibitor of MerTK (10nM, UNC2025, MedChemExpress, Monmouth Junction, NJ). MOVAS cells were labeled with PKH67 green-fluorescent dye (Sigma-Aldrich, St. Louis, MO). Labeled cells were then treated with 1 µM staurosporine (Thermo Fisher Scientific, Waltham, MA) for a total of 2 hours at 37°C to induce apoptosis. Apoptotic MOVAS cells were then washed and co-cultured with isolated peritoneal macrophages at a ratio of 1:1 (1 x 10^6^ cells each) in 6-well plates. These were incubated together at 37°C for 45 minutes and washed with PBS. Cells were then stained for Live/Dead staining (Live/Dead Blue, Thermo Fisher Scientific, Waltham, MA), then subsequently stained with PE-Cy7-conjugated anti-mouse F4/80 antibody (BioLegend, San Diego, CA) to localize macrophages. Efferocytosis was quantified through flow cytometry, using the percentage of F4/80^+^ macrophages that were synchronously positive for PKH67 fluorescence, thus signifying engulfment of apoptotic MOVAS cells.^87^

Following the 45-minute efferocytosis assay, non-engulfed apoptotic MOVAS cells were removed by three gentle washes with ice-cold PBS. Macrophages were subsequently incubated for 4 hours at 37°C in fresh RPMI-1640 supplemented with 0.5% FBS. To maintain sustained MerTK inhibition throughout the post-efferocytosis incubation period, UNC2025 (10 nM) was included in the fresh medium for the UNC2025 and Evolocumab+UNC2025 treatment groups. Following the 4-hour incubation, culture supernatants were collected and stored at −80°C. Adherent macrophages were washed twice with ice-cold PBS and lysed by addition of [150 µL] RIPA supplemented with 1x protease and phosphatase inhibitor cocktail per well. Plates were incubated on ice for 20 minutes with gentle rocking, after which cells were scraped and transferred to pre-chilled 1.5 mL microcentrifuge tubes. Lysates were clarified by centrifugation at 12,000xg for 15 minutes at 4°C, and supernatants were collected and stored at −80°C until analysis. Total protein concentration was determined using the Pierce BCA Protein Assay Kit (Thermo Scientific, Waltham, MA) per the manufacturer’s instructions, and all lysate samples were normalized to equal protein concentrations prior to ELISA. Intracellular arginase1 (ARG1) protein levels were quantified using the Mouse ARG1 Rapid ELISA Kit (Thermo Fisher Scientific, Waltham, MA; cat. No EELR124) according to the manufacturer’s protocol.

### Statistical analysis

Statistical analyses were performed on GraphPad Prism 10 (GraphPad Software, La Jolla, CA) software. For comparisons between three or more cohorts, a one-way ANOVA followed by Tukey’s multiple comparisons test was performed. Either unpaired Student’s *t test* or a non-parametric Mann-Whitney test were performed for pairwise comparisons. Data are presented as mean ± standard error of the mean. For the retrospective clinical component, raw and matched comparisons were performed using Kruskal-Wallis (continuous variables) and Fisher’s exact tests (categorical variables). Kaplan-Meier estimates were used to determine long-term survival estimates based on therapy. For survival analysis, the time between initial and most recent AAA measurement was taken as a surrogate survival time for surviving patients. Continuous variables are presented as median [IQR] (range); categorical variables presented as N(col%). Statistical significance was defined as p<0.05. Sparse partial least squares discriminant analysis (sPLS-DA) and Variable Importance in Projection (VIP) plots of lipidomic and metabolomic data were generated using the MetaboAnalyst platform. Prior to analysis, the raw data were normalized, log-transformed, and auto-scaled to improve comparability across samples. VIP scores were subsequently generated from the PLS-DA model to rank metabolites based on their contribution to group discrimination.

## Notes

### Competing Interest Statement

The authors have declared no competing interest.

