## Supplemental Figures 1-4 for "Proprotein convertase subtilisin kexin type 9 (PCSK9) inhibition attenuates abdominal aortic aneurysm formation via enhanced macrophage-dependent efferocytosis"

S1

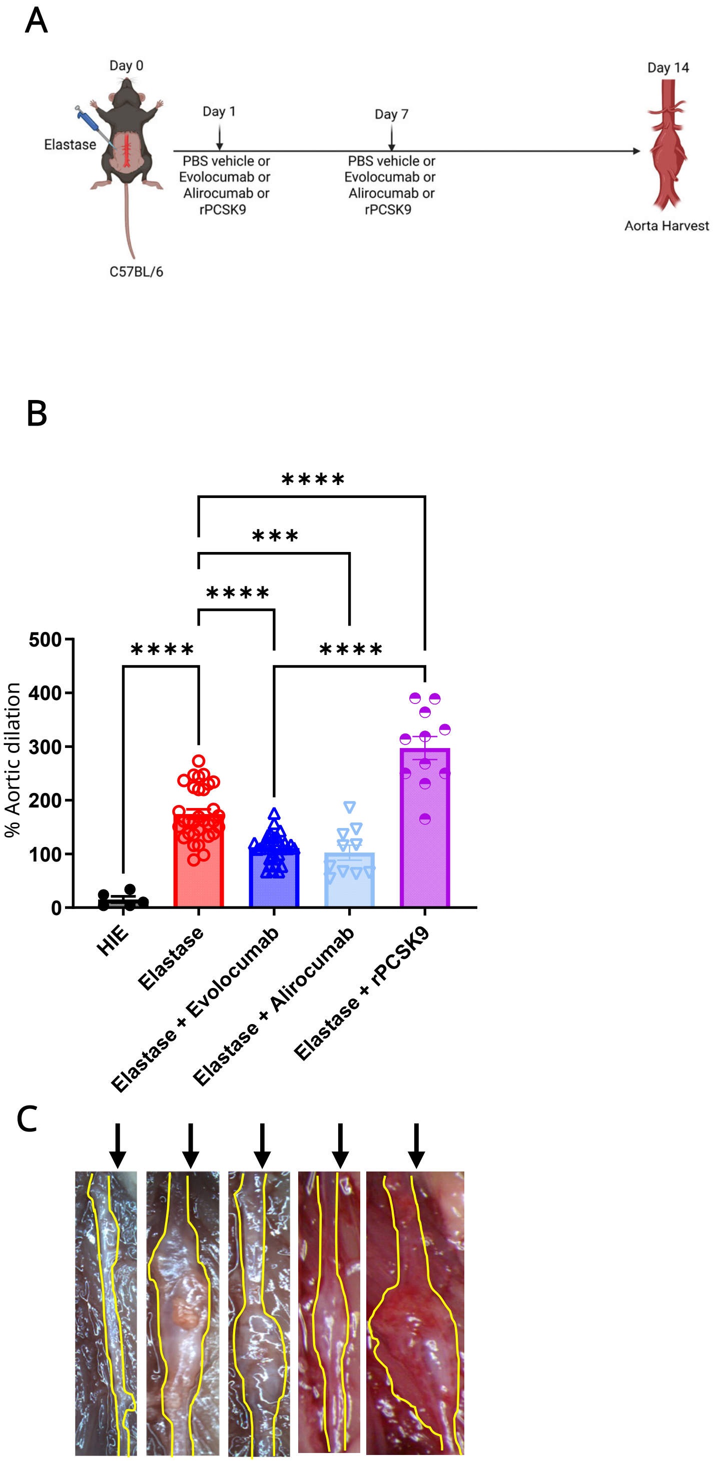


**S1 | AAA growth is accentuated with recombinant PCSK9 treatment and attenuated with exposure to monoclonal PCSK9 inhibitor therapies. a**, Schematic representation of topical elastase model with intraperitoneal injection of PBS control vehicle, Evolocumab (10 mg/kg), Alirocumab (10 mg/kg) or recombinant PCSK9 (0.2 µg/µL) on days 1 and 7. **b**, Mice treated with elastase and recombinant PCSK9 exhibited significant increase in AAA diameter compared to elastase alone and control treated mice. Conversely, treatment with either Evolocumab or Alirocumab resulted in comparable reductions in AAA growth compared to the elastase alone treated mice. n=5-32/group. **p<0.005, ***p<0.0005, ****p<0.0001, ns=not significant. **c**, Representative images of aortic growth at the time of harvest.

S2

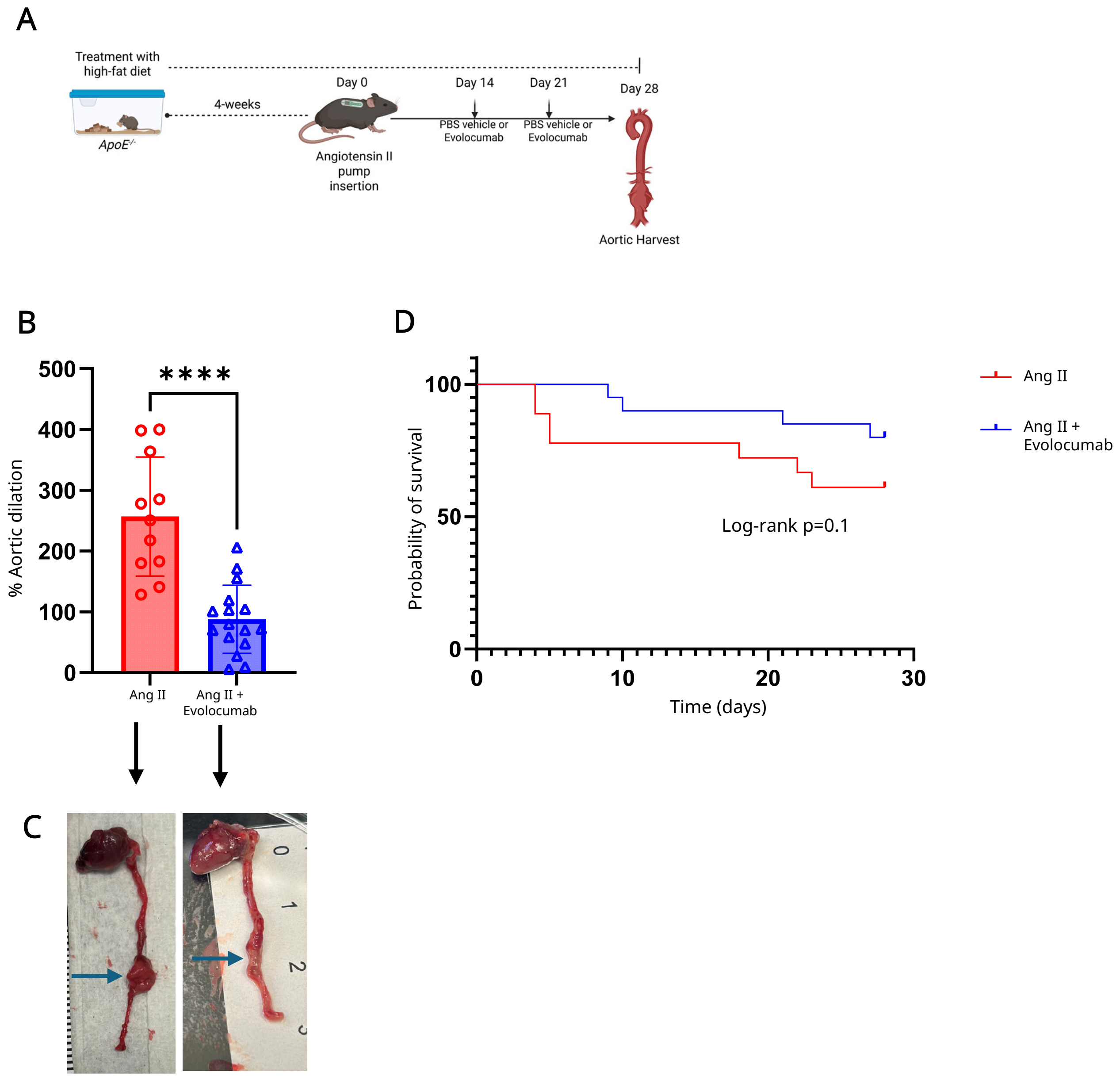


**S2|PCSK9 inhibition attenuates aneurysm growth in Angiotensin-II/ApoE$^{-/-}$ rupture model. a**, Schematic representation of Angiotensin II/*ApoE$^{-/-}$* model with intraperitoneal injection of PBS vehicle or Evolocumab on days 14 and 21. **b**, Evolocumab-treated mice significantly attenuated aortic growth compared to untreated mice. n=11-16/group; ****p<0.0001. **c**, Representative imaging of aortic diameters on day 28. **d**, Kaplan Mier survival estimates from rupture-related mortality (Angiotensin II/*ApoE$^{-/-}$* 61% vs. Angiotensin II/*ApoE$^{-/-}$* + Evolocumab 80%; log-rank p=0.1).

S3

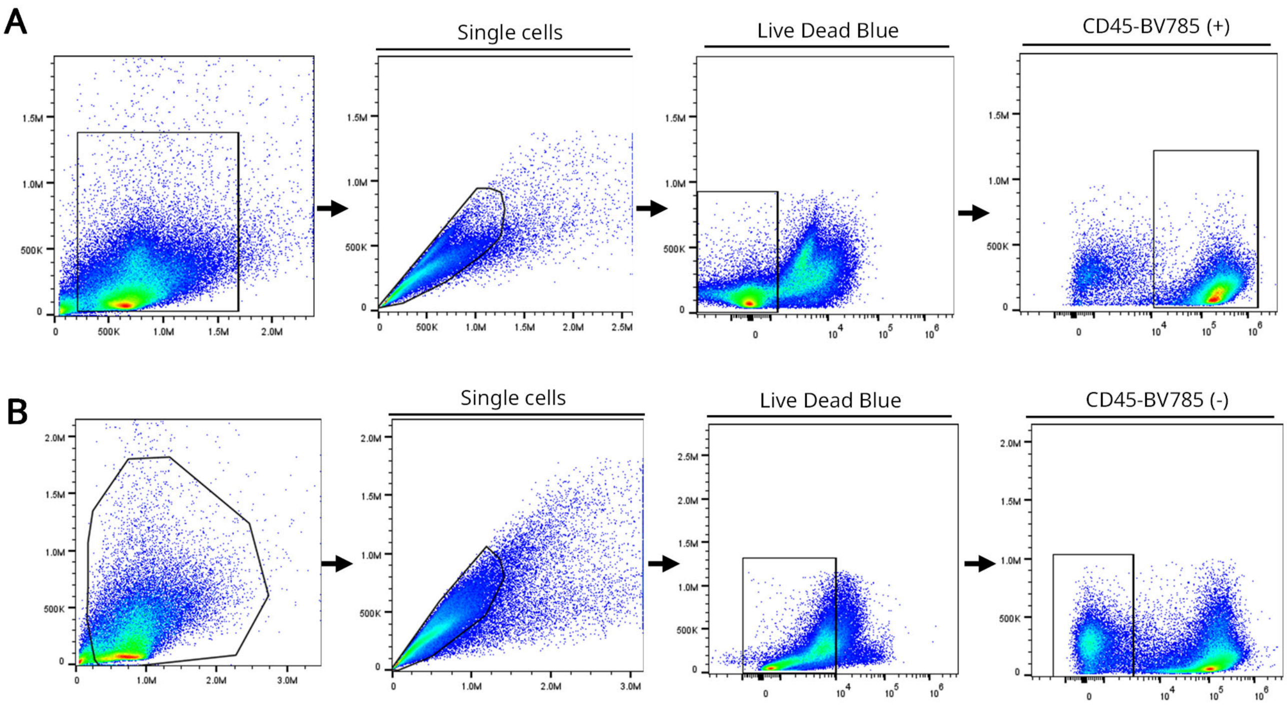


**S3|Flow cytometry gating strategies for *in vivo* efferocytosis assay. a**, Schematic representation of the gating strategy used for flow cytometry analysis of efferocytosis using *in vivo* aortic tissue from elastase-treated mice with or without Evolocumab treatment. **b**, Gating strategy of cleaved-caspase 3 assay for *in vivo* flow cytometry analysis in elastase-treated mice with or without Evolocumab treatment.

S4

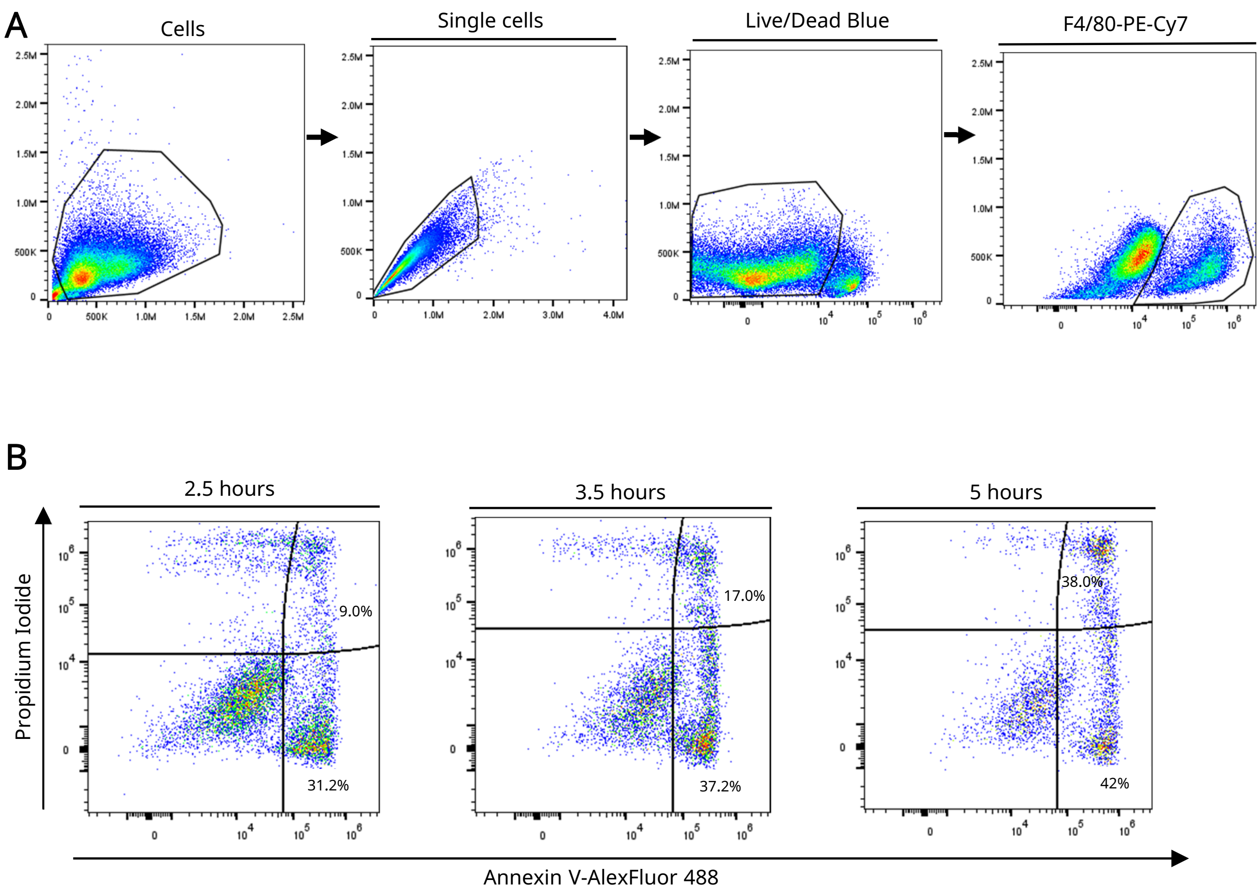


**S4 | Flow cytometry gating strategies for *in vitro* efferocytosis assay. a**, Schematic representation of the gating strategy used for flow cytometry analysis of *in vitro* efferocytosis in cultures of macrophages and SMCs (MOVAS cells). **b**, MOVAS apoptosis assay in which cells were treated with 1 μM staurosporine for varying time intervals and apoptotic cells were quantified using Annexin-V and propidium iodide staining.
